# Subcapsular sinus macrophage sensing of extracellular matrix rigidity alters membrane topography and immune complex mobility

**DOI:** 10.1101/2022.12.02.518873

**Authors:** Maro Iliopoulou, Anna T. Bajur, Hannah C. W. McArthur, Carl Coyle, Favour Ajao, Robert Köchl, Andrew P. Cope, Katelyn M. Spillane

## Abstract

Subcapsular sinus macrophages (SSMs) play a key role in immune defence by forming immunological barriers that control the transport of pathogens from lymph into lymph node follicles. SSMs participate in antibody responses by presenting antigens directly to naive B cells and by supplying antigens to follicular dendritic cells to propagate germinal centre reactions. Despite the prominent roles that SSMs play during immune responses, little is known about their cell biology because they are technically challenging to isolate and study *in vitro*. Here, we used multi-colour fluorescence microscopy to identify lymph node-derived SSMs in culture. We focused on the role of SSMs as antigen-presenting cells and found that their actin cytoskeleton regulates the spatial organisation and mobility of immune complexes displayed on the cell surface. Moreover, we determined that SSMs are mechanosensitive cells that respond to changes in extracellular matrix (ECM) rigidity by altering the architecture of the actin cytoskeleton, leading to changes in cell morphology, membrane topography, and immune complex mobility. Our results reveal a new mechanism regulating physical aspects of antigen presentation by antigen-presenting cells, which may have implications for B cell activation and antibody responses.

## Introduction

Adaptive immune responses rely on the arrival of exogenous antigens to lymph node follicles, where they can trigger B cell activation to initiate antibody responses. Antigens arrive in lymph nodes via afferent lymphatic vessels, which propel lymph-borne antigens over a layer of CD169^+^ macrophages that line the subcapsular sinus floor. Subcapsular sinus macrophages (SSMs) have a distinct morphology that underpins their dual roles as immunological barriers and antigen-presenting cells. They interdigitate a layer of lymphatic endothelial cells (1), protruding a “head” into the lymph to capture antigens (2–5), and extending long “tail” processes deep into the follicle either to present antigens directly to cognate B cells (4) or to transfer them to follicular dendritic cells (FDCs) for long-term retention (6–8). SSMs thus play a key role in naive B cell activation by acting as antigen-presenting cells, and in B cell affinity maturation by controlling antigen deposition on FDCs, which supply antigens to B cells during germinal centre reactions.

The prominent role that SSMs play in B cell responses has prompted several groups to attempt to isolate SSMs by lymph node digestion and cell sorting to study their cell biology (3, 5, 6, 9–11). However, SSMs are limited in number, highly sensitive to manipulation, and prone to die during isolation protocols. They form apoptotic membrane blebs that are acquired by interacting lymphocytes, which then carry SSM surface markers and masquerade as SSMs during flow cytometric analysis (12). SSMs are, therefore, technically challenging to study *in vitro*, and consequently we have limited understanding of their biology at the single-cell level.

Here, we have used large-scale, multi-colour fluorescence imaging to identify SSMs in in *vitro* mouse lymph node cultures. We demonstrate that the SSM actin cytoskeleton regulates both the spatial organisation and mobility of immune complexes presented on the cell surface, and that these physical features of antigen presentation are altered by changes to ECM rigidity.

## Results

### Identifying SSMs *in vitro*

Identification of SSMs in culture experiments was fortuitous and arose through studies of antigen presentation by FDCs (13). In these experiments we extract FDCs from mouse lymph nodes through a combination of mechanical and enzymatic disruption of tissue followed by positive selection using antibody complexes containing an FDC-specific antibody (rat IgG2c, *κ* anti-mouse FDC-M1). This approach enriches FDCs but does not purify them, and FDCs must be identified in culture as CD45^-^ CD21/35^+^ cells that capture and retain immune complexes on their surface (14–16). During our investigations, we identified unexpectedly a population of CD45^+^ CD21/35^-^ cells that were also heavily labelled by immune complexes, and so used multi-colour fluorescence imaging and flow cytometry to immunophenotype the cells in more detail, based on surface marker expression.

Flow cytometric analysis showed that the FDC-M1 positive selection enriched CD45^+^ CD11b^+^ cells of myeloid lineage (Fig S1, A and B). These cells are known to differentiate into either macrophages or dendritic cells and populate lymphoid organs (17). Lymph node macrophages and dendritic cells can be distinguished based on expression of CD11c; dendritic cells are CD11c^hi^ while macrophages are CD11c^neg/low^ (6, 18). We observed by flow cytometry that >80% of CD11b^+^ cells were CD11c^neg/low^ (Fig S1B), and fluorescence imaging of the cells stained with anti-CD11b and anti-CD11c monoclonal antibodies in culture confirmed that they expressed CD11b (Fig. 1A) but not CD11c (Fig. 1B). This CD45^+^ CD11b^+^ CD11c^neg/low^ phenotype indicated that the cells were likely macrophages. We confirmed further their macrophage lineage by staining them with an antibody targeting CD68, a transmembrane glycoprotein that is highly expressed by macrophages and localises primarily to endosomes and lysosomes (Fig. 1C) (19).

**Fig. 1:**
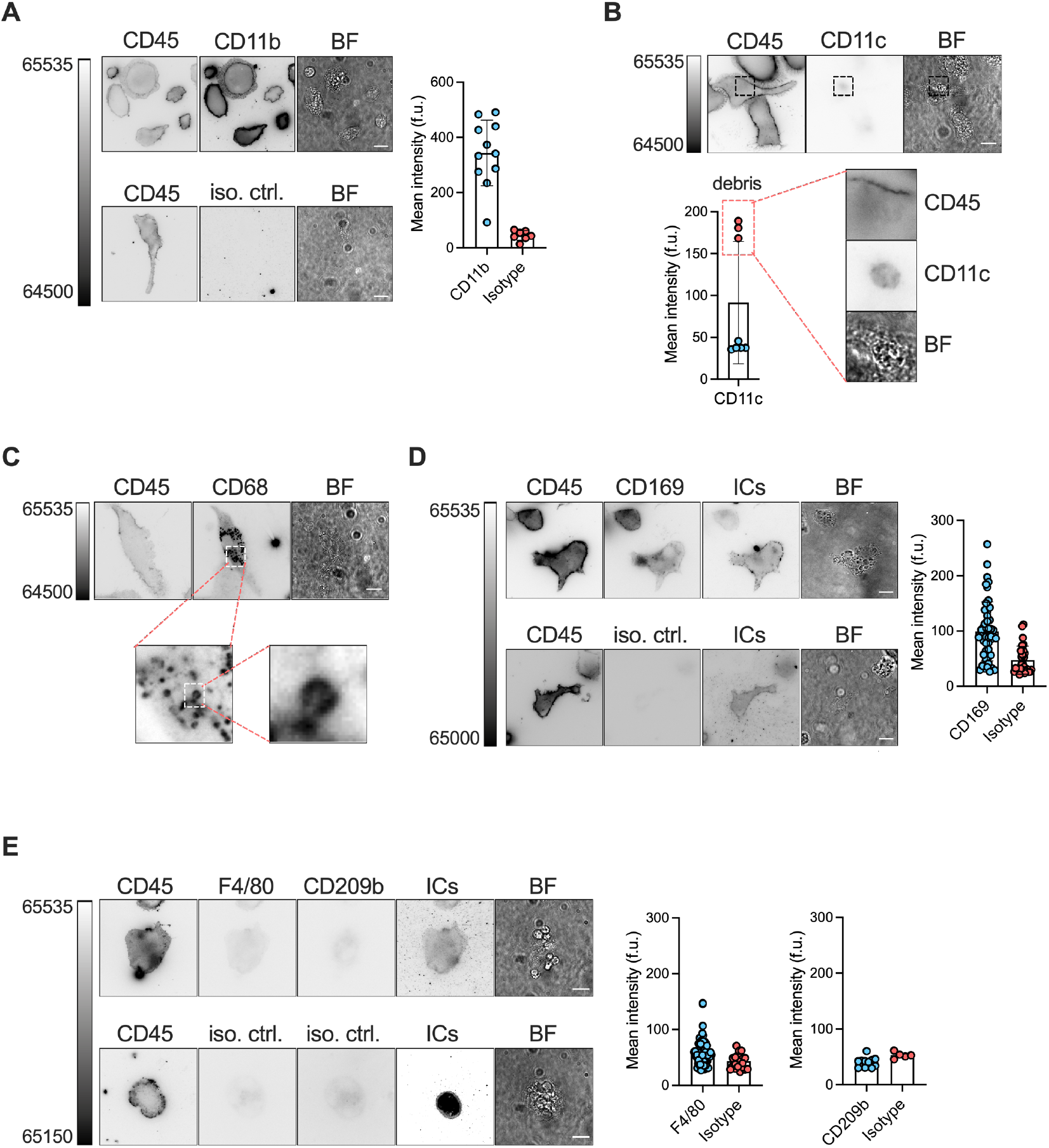
Identification of SSMs in culture with multi-colour fluorescence microscopy. SSMs were enriched from lymph node single-cell suspensions, cultured *in vitro* on collagen I-coated class coverslips, labelled with fluorescent immune complexes (ICs), and stained with monoclonal antibodies to CD45, CD11b, CD11c, CD68, CD169, F4/80, and CD209b. Immunofluorescence imaging and quantitation confirmed that the cells are (A) CD45^+^ CD11b^+^, (B) CD11c^-^, (C) CD68^+^, and (D) CD169^+^. As expected, antibodies against CD45, CD11b, and CD169 specifically label the plasma membrane, while anti-CD68 labels the membranes of intracellular vesicles. Though several cells had relatively high mean intensity values of CD11c staining, close inspection of the images reveals that this signal is due to cell debris and not specific staining of the cell membrane. (E) The cells expressed no or very low amounts of the medullary macrophage markers F4/80 and CD209b. n = 5 to 48 SSMs collated from 2 biological replicates. Scale bars: 10 μm.

Lymph node macrophages are a heterogeneous population of cells that, during homeostasis, are classified into three subsets: subcapsular sinus macrophages (SSMs), medullary sinus macrophages (MSMs), and medullary cord macrophages (MCMs) (12, 20). These subsets can be distinguished based on their surface expression of the C-type lectin sialoadhesin (CD169) and the murine macrophage marker F4/80. MCMs are CD169^-^ while SSMs and MSMs are both CD169^+^. The latter two populations differ in F4/80 expression; SSMs are CD169^+^ F4/80^neg/low^ and MSMs are CD169^+^ F4/80^+^ (21). Immunofluorescence staining of cells cultured on collagen I-coated glass showed that they expressed CD169 (Fig. 1D) and were negative or expressed very low levels of F4/80 (Fig. 1E), while flow cytometric analysis confirmed that >90% of CD169^+^ cells were F4/80^-^ (Fig S1B). Immunofluorescence staining also showed that the cells were negative for SIGN-R1 (CD209b) (Fig. 1E), a marker of medullary macrophages not expressed by mouse SSMs (Fig. 1E) (12, 21). Based on this extended CD11b^+^ CD11c^-^ CD169^+^ F4/80^neg/low^ CD209b^-^ phenotype and the ability of the cells to capture and retain large amounts of immune complexes on their surfaces, we concluded that the cells are indeed SSMs.

### SSMs use Fc*γ* receptors to present immune complexes

SSMs do not express FDC-M1 but do express high levels of Fc*γ* receptors (Fc*γ*Rs) (22). We therefore postulated that SSMs were enriched during the cell isolation procedure due to interactions between their FcγRs and the Fc domain of either the rat IgG2c, *κ* anti-mouse FDC-M1 primary antibody or the biotinylated mouse IgG2a anti-rat Ig*κ* secondary antibody used for cell enrichment. First, we confirmed that cells expressed FcγRs by staining with anti-CD16/32 (Fc*γ*RIII/Fc*γ*RΠ) (Fig. S2A). Next, we repeated the cell isolation using a rat IgG2c, *κ* isotype control antibody in place of the FDC-M1 antibody. Flow cytometric analysis confirmed our hypothesis that SSMs were enriched through interactions between IgG antibody complexes and Fc*γ*Rs (Fig. S1C). Given the high capacity of SSMs to bind IgG antibody complexes, we wondered whether these cells require complement to present immune complexes. To answer this question, we loaded SSMs with antibody complexes composed of goat IgG anti-mouse *κ* and donkey anti-goat IgG (H+L) that either were, or were not, mixed with serum in GVB++ buffer as a source of complement. We observed that complement fixation was not required for SSMs to capture and retain antibody complexes on the cell surface (Fig S2, B and C). Therefore, we conclude that SSMs primarily use Fc*γ*Rs to present immune complexes.

### Immune complexes on the SSM surface colocalise with actin filaments

We next characterised the morphology of SSMs at the single-cell level *in vitro*. After culturing SSMs on collagen I-coated glass coverslips for 5 to 7 days, we loaded them with Cy3B-labelled immune complexes and then fixed, permeabilised, and stained them with phalloidin-Alexa Fluor 488 (AF488) to detect filamentous (F)-actin (Fig. 2A). Staining the actin cytoskeleton revealed that SSMs form several distinct actin structures including filopodia (Fig. 2B), lamellipodia (Fig. 2C), and podosomes (Fig. 2D) at the cell-ECM interface and actin-enriched membrane ruffles (Fig. 2E) on the dorsal cell surface. Immune complexes were excluded from podosomes but colocalised with both filopodia and ruffles, and heavily decorated lamellipodia. We quantified the spatial association of immune complexes with Factin, focusing on membrane ruffles. To do this, we developed an image analysis workflow that assigned each immune complex on the dorsal cell surface as either “on actin” or “off actin”, to determine the fraction of immune complexes that associate with ruffles (see materials and methods). The quantitation showed that all immune complexes on the dorsal cell surface associated with membrane ruffles (Fig. 2, E and F). Treatment of the cells with pharmacological inhibitors of proteins that regulate actin assembly or contractility, including CK-666 (Arp2/3), SMIFH2 (formins), blebbistatin (myosin II), and Y-27632 (Rho-associated protein kinase), slightly loosened the association of immune complexes with actin ruffles but did not disrupt it (Fig. 2, F and G; Fig. S3). Treating cells with jasplakinolide to stabilise actin filaments slightly improved the spatial overlap of immune complexes and ruffles. Only mycalolide B, a drug that severs actin filaments, completely disrupted the association by removing dorsal membrane ruffles from the cells (Fig. S3). We also confirmed that immune complex association with cytoskeletal filaments was specific to F-actin, as they did not colocalise with microtubules or the intermediate filaments vimentin and nestin (Fig. 2H; Fig. S4, A–C). These results demonstrate that immune complexes have a strong propensity to localise to actin-enriched membrane ruffles on the dorsal surfaces of SSMs.

**Fig. 2:**
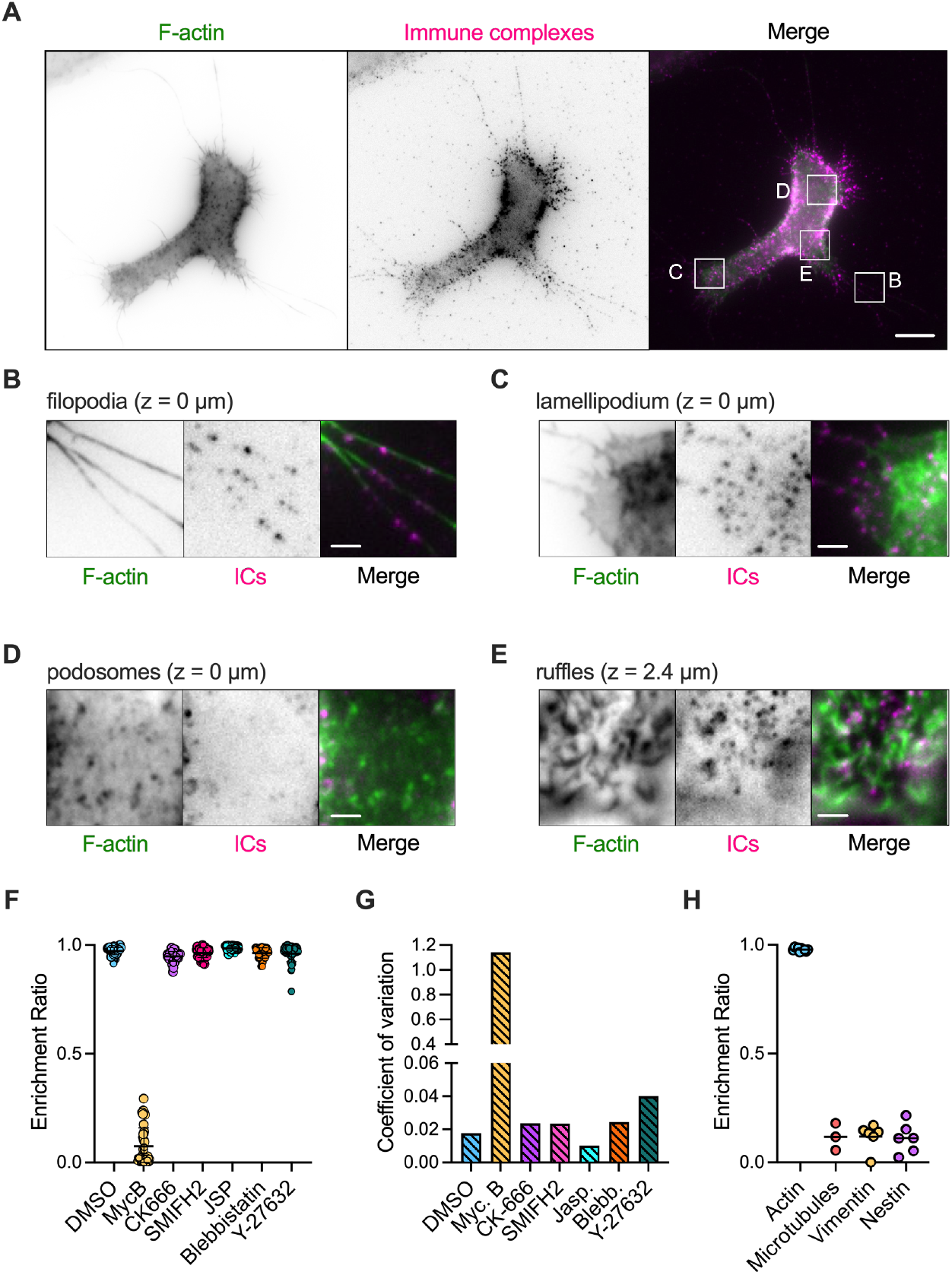
Immune complexes presented by SSMs associate with actin filaments. (A) A single z-plane image (z = 0 μm) from a z-stack of an SSM labelled with immune complexes and stained with phalloidin-AF488. Scale bar: 10 μm. (B-E) Zoomed in regions highlighted in (A) showing the different F-actin structures formed by SSMs including (B) filopodia, (C) lamellipodia, (D) podosomes, and (E) dorsal membrane ruffles. Immune complexes densely label filopodia, lamellipodia, and ruffles and are excluded from podosomes. Scale bars: 2 μm. (F) Quantitation of the spatial association of immune complexes with dorsal membrane ruffles in SSMs treated with drugs that perturb actin filament assembly or actomyosin contractility. An enrichment ratio value of 1 indicates “on actin” and a value of 0 indicates “off actin.” Each dot represents the mean enrichment ratio value of all immune complexes within six regions of interest (ROIs) from one cell. n = 44-57 cells per treatment, pooled from six independent experiments. (G) Coefficient of variation values for the data in 2F. (H) Quantitation showing that immune complexes associate with F-actin-rich dorsal membrane ruffles, but not microtubles, vimentin, or nestin.

### Immune complex mobility coincides with F-actin dynamics

The actin cytoskeleton regulates the functions of immunoreceptors in many immune cell types including phagocytic macrophages, where binding of IgG antibodies to Fc*γ*Rs induces cytoskeletal remodelling and consequent changes in Fc*γ*R mobility and clustering (23). Since we observed that F-actin tightly associates with Fc*γ*R-presented immune complexes in fixed SSMs, we reasoned that the cell cytoskeleton might also constrain the lateral mobility and clustering of immune complexes in live SSMs. To investigate this possibility, we analysed the motion of fluorescent immune complexes on the dorsal surfaces of SSMs expressing LifeAct-green fluorescent protein (GFP) (Fig. 3A), which binds specifically to F-actin in live cells without interfering with actin dynamics (24). Immune complex labelling was performed at low density so that individual immune complexes could be detected. The cell surface was imaged in the immune complex and F-actin channels with a frame rate of 20 Hz for 15 seconds using widefield epifluorescence microscopy at 37 °C (Movie S1).

**Fig. 3:**
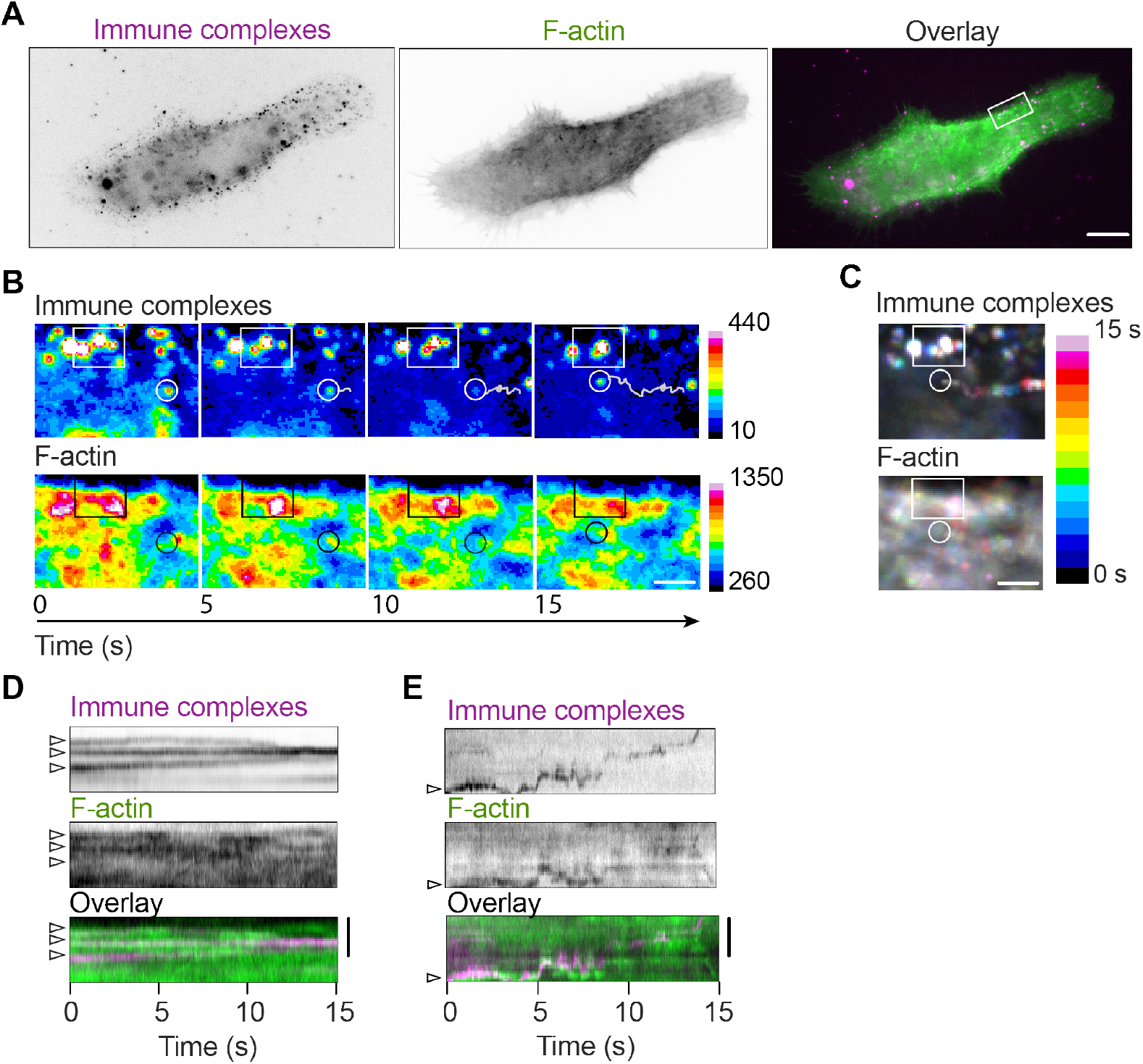
The movement of F-actin and immune complexes are spatiotemporally correlated at the SSM surface. (A) Representative widefield image of F-actin (LifeAct-GFP, green) and immune complexes (Cy3B-labelled, magenta) at the dorsal membrane of a live SSM. Scale bar: 10 μm. (B) Magnification of the boxed area in (A). Two-colour time-lapse images of immune complexes and F-actin. Images are pseudo-coloured to represent changes in fluorescence intensity. Low-mobility immune complexes are denoted by a white box and by the corresponding black box in the F-actin images. These complexes were trapped in a region of high F-actin intensity. A high-mobility immune complex is denoted by a white circle and the corresponding black circle in the F-actin images. The trajectory traces the movement of the mobile immune complex across time. The high F-actin intensity immediately adjacent to the complex indicates that the complex is “pushed” by a dynamic F-actin structure. Scale bar: 2 μm. (C) Temporal projections of immune complex and F-actin dynamics (t = 0 to 15 s) with cold colours representing early times and warm colours later times. Scale bar: 2 μm. (D, E) Kymographs representing the motion of immune complexes and F-actin from (D) the boxed region and (E) along the trajectory marked in (B). Arrow heads show the starting positions of immune complexes in each kymograph. Vertical scale bars: 2 μm.

In agreement with the fixed-cell images, the live-cell timelapse images showed that immune complexes accumulated near regions of high F-actin intensity (Fig. 3B). The motion of immune complexes and F-actin structures was correlated; slow-moving immune complexes associated with large, static F-actin aggregates while highly mobile immune complexes appeared to be “pushed” by small, dynamic F-actin patches (Fig. 3, B and C, and Supplementary Video 1). The coincidence of immune complex and F-actin dynamics was further supported by kymographic analysis. Within dense regions of F-actin, low-mobility immune complexes were brought together to form larger clusters, indicating that F-actin can induce clustering of immune complexes on the cell surface (Fig. 3D). Elsewhere on the cell membrane, fast-moving immune complexes and F-actin patches showed that they were spatiotemporally linked (Fig. 3E). Taken together, these observations indicate that the SSM actin cytoskeleton regulates the organisation and mobility of immune complexes on the cell surface.

### Immune complexes presented by SSMs exhibit multiple diffusive states

The previous results showed that immune complexes spatially and temporally associate with Factin in SSMs expressing LifeAct-GFP. To explore further the role that F-actin plays in controlling immune complex motion, we characterised the diffusion of immune complexes displayed by cells that had been treated with mycalolide B to depolymerise actin filaments, jasplakinolide to stabilise actin filaments, or DMSO as a control. Visual inspection of the immune complex trajectories with and without drug treatments indicated that the SSM actin cytoskeleton did influence immune complex mobility (Fig. 4A). In control (DMSO-treated) cells, we observed immobile and partly mobile populations as before (Fig. 3). Treatment with mycalolide B visibly increased immune complex mobility (Fig. 4A, Movie S2) and treatment with jasplakinolide immobilised all immune complexes (Fig. 4A, Movie S3).

**Fig. 4:**
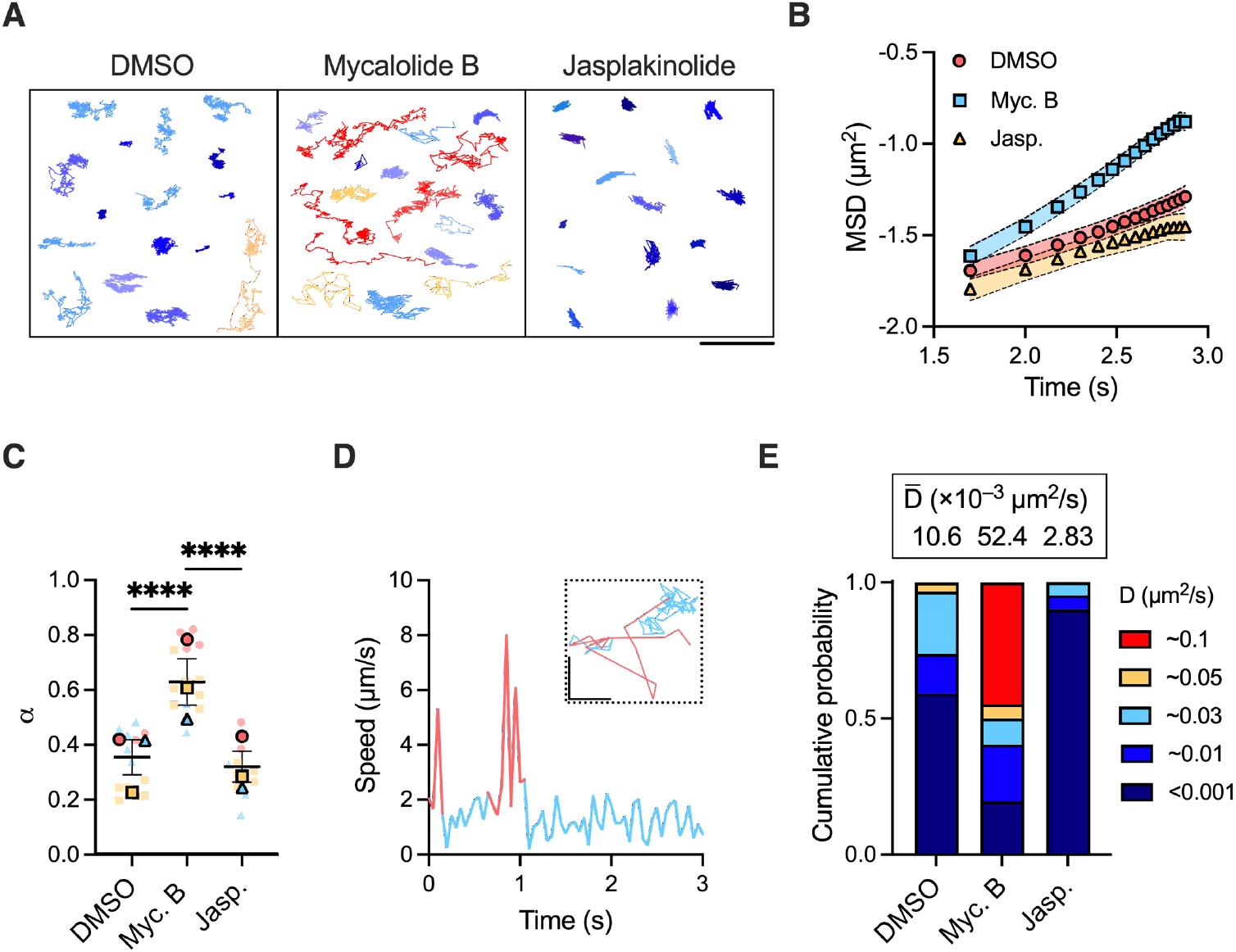
Quantitative analysis of immune complex single-particle trajectories. (A) Representative trajectories of individual immune complexes diffusing on the surfaces of SSMs treated with DMSO, mycalolide B, or jasplakinolide. Trajectories are colour-coded by their diffusion constant, with highly mobile complexes in red (D≈0.1 μm^2^/s) and immobile complexes in dark blue (D<0.001 μm^2^/s). Scale bar: 200 nm. (B) Log-log plot of MSD versus time from immune complex single-particle trajectories. The symbols represent the mean MSD values from all acquired trajectories per condition (>1000 trajectories per cell; DMSO: 13 cells, Myc B: 14 cells, Jasp.: 10 cells), and the fills the standard errors of the mean. Data are from one experiment representative of three experiments. (C) Anomalous scaling exponents, *α*, extracted as the slopes from linear fits to plots of MSD versus time for the first 10 time lags. Each plain dot represents the mean *α* value for >1000 immune complexes from one cell (n = 2-7 cells per condition) and each outlined dot represents the mean value for one independent experiment (N = 3 experiments). Data are colour coded by experiment. Bars represent mean ± SEM. (D) Sample plot of instantaneous speed versus time for one immune complex. Instances of high speed are colour-coded pink and instances of low speed are blue. The corresponding xy trajectory is shown in the inset with the same colour-coding scheme. Inset scale bars: 200 nm. (E) Bar graphs showing the mean weight fraction, *π*, of each mobility state identified by SMAUG analysis for immune complexes diffusing on cells treated with DMSO, mycalolide B, and jasplakinolide. The plot was constructed from the same single-particle trajectories used to calculate the values of α in (C) and merges data from three independent experiments. The weighted mean of the diffusion constant for each condition, 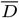, is given atop each bar. ****p<0.0001.

The diffusion of a molecule can be categorised as one of three types based upon the relationship between its mean-squared displacement (MSD), 〈r^2^ (*τ*)〉, and time lag, *τ*: Brownian (random), superdiffusive (directed), and subdiffusive (confined). The MSD of particles diffusing in the plane of a membrane is described by the power law 〈r^2^ (*τ*)〉 = 4D*τ^α^*, where D is the diffusion constant and *α* is the anomalous scaling exponent. Values of *α* = 1 indicate Brownian motion, *α* < 1 subdiffusive motion, and *α* > 1 superdiffusive motion. A log-log plot of MSD versus *τ* yields a straight line of slope *α* and thus provides a convenient representation of the nature of the motion (Fig. 4B). Our results show that immune complex diffusion was confined in control cells (*α* = 0.38) (Fig. 4C), in agreement with observations that Fc*γ*Rs are confined within submicron actin compartments in bone marrow-derived macrophages (23). Treatment of SSMs with jasplakinolide to stabilise actin filaments slightly increased immune complex confinement (*α* = 0.35) while treatment with mycalolide B to remove actin barriers drastically reduced confinement (*α* = 0.62) (Fig. 4C). Thus, our data indicate that the actin cytoskeleton constrains the lateral mobility of immune complexes on the SSM surface.

MSD analysis provides a single diffusion constant for each immune complex. Our observations that immune complexes could transiently associate with actin, however, suggested that each immune complex could sample different mobility states within a single trajectory (Fig. 3E and 4A). This observation was supported by plotting the instantaneous velocity of immune complexes over time, which revealed that they generally moved with slow speeds of <2 μm/s but were transiently free to move with speeds of 5-8 μm/s (Fig. 4D). To quantify this heterogeneous behaviour and better understand the role of actin in generating distinct mobility characteristics, we analysed the single-particle trajectories using single-molecule analysis by unsupervised Gibbs sampling (SMAUG) (25). SMAUG uses nonparametric Bayesian statistics to uncover different mobility states from within single trajectories and quantify their diffusion constants (D) and weight fractions (π). We applied this analysis tool to our trajectories and found that in DMSO-treated cells, most immune complexes were immobile (π_immobile_=0.6, D_immobile_<0.001 μm^2^/s), with the remaining having moderate mobility (π_moderate_=0.4, Dmoderate=0.01 to 0.05 μm^2^/s) (Fig. 4E). Treatment with mycalolide B to sever actin filaments allowed immune complexes to diffuse more freely. While the fraction of complexes with moderate mobility was slightly lower (π_moderate_=0.35), the proportion of immobile complexes was substantially reduced (πimmobile=0.2) and a new, highly mobile population appeared (π_high_=0.45, D_high_=0.1 μm^2^/s). Treatment with jasplakinolide to stabilise actin filaments caused almost complete immobilisation of immune complexes (π_immobile_=0.9). Taken together, these data indicate that the SSM actin cytoskeleton constrains the mobility of immune complexes presented by Fc*γ*Rs on the cell surface.

### ECM rigidity alters SSM morphology

Our results suggested that dorsal membrane ruffles are morphological structures involved in immune complex presentation by SSMs. Generally, the morphology of a cell results from the balance of cell-intrinsic forces such as tension and contractility, and cell-extrinsic forces such as ECM stiffness (26). Disrupting the cellular force balance can cause rapid alterations in cy-toskeletal structure (27) that can impact cell polarisation (28) and membrane ruffling (29). We were therefore curious to know whether SSMs could also alter the organisation of their actin cytoskeleton to reflect changes in ECM rigidity.

To investigate the effect of ECM stiffness on SSMs, we cultured the cells on glass and flat polyacrylamide hydrogels with similar surface densities of collagen I (Fig. S5A) but different rigidities, with mean Young’s moduli ranging from 2 to 180 kPa (Fig. 5A; Supplementary Table 3). SSMs cultured on the gels for 5 days adhered well to all of the gels (Fig. 5B). Imaging F-actin revealed that compared to cells cultured on glass, cells cultured on 2 kPa gels had a 60% lower mean spread area (Fig. 5C), 43% increased mean roundness (Fig. 5D), and 61% lower cell volume (Fig. 5E). These results indicate that SSMs become smaller and more rounded in response to a softer ECM.

**Fig. 5:**
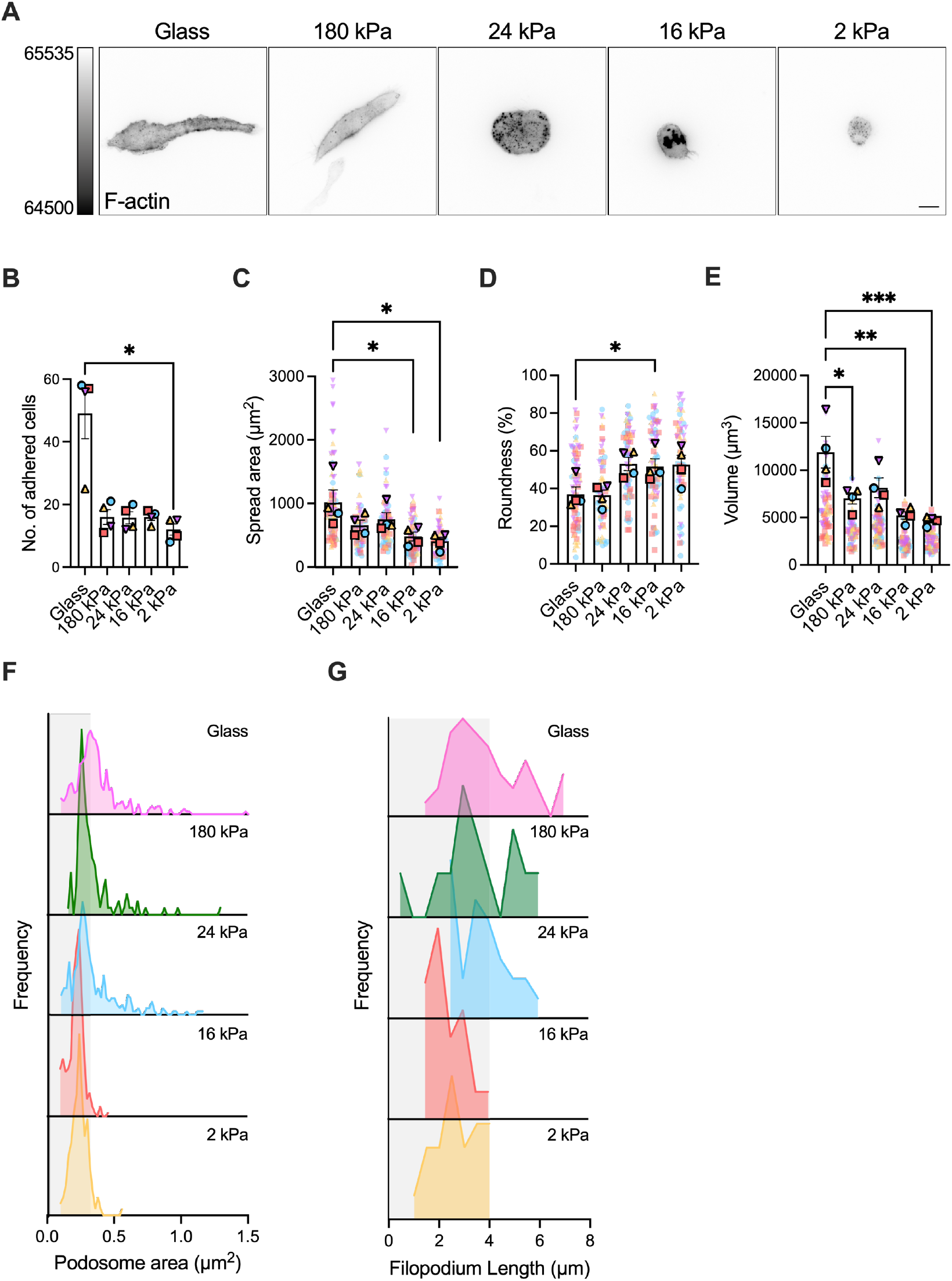
ECM rigidity alters SSM morphology and actin cytoskeleton organisation. (A) Images of SSMs on collagen I-coated substrates of different stiffness. Scale bar: 10 μm. (B) The number of SSMs adhered to the 78.5 mm^2^ substrate surface on day 5 of culture. Data are from four independent experiments. (C, D) Cells change their size and shape in response to ECM rigidity as assessed by (C) spread area (μm^2^), (D) roundness (%), and (E) volume (μm^3^). Each plain dot represents one cell and each outlined dot represents the mean value for one independent experiment (n = 11 to 26 cells per condition per experiment, N = 4 independent experiments). Data are colour-coded by experiment. Bars represent mean ± SEM. (F-H) Cells respond to changes in substrate stiffness by altering (F) the size of podosomes at the ECM-cell interface and (G) the length of filopodia. The bin sizes for the histograms in (F) and (G) are 0.02 μm^2^ and 0.5 μm, respectively. Data are pooled from three or four independent experiments (n = 15 to 34 cells per condition per experiment). *p<0.05, **p<0.01, ***p<0.001.

SSM responses to ECM rigidity were also reflected by changes in the prominence of F-actin-based structures. Podosomes are actin-rich, circular structures that form at the ventral surface of dendritic cells and macrophages, enabling cells to adhere to the ECM via integrins (30). They also have a mechanosensory function, allowing cells to sense the rigidity of the substrate. SSMs on all substrates formed podosomes. Reducing ECM stiffness did not cause a major change in the mean area of the podosome actin core (0.29 μm^2^ on glass and 0.22 μm^2^ on 2 kPa gels) (Fig. S5B), but detailed analysis showed that the number of podosomes with area >0.3 μm^2^ decreased substantially on soft ECM (41% on glass versus 15% on 2 kPa gels) (Fig. 5F). Likewise, filopodia—thin membrane protrusions that cells use to probe the microenvironment—were formed by SSMs on all substrates but grew shorter on soft ECM (Figs. 5G and S5C), with the proportion of filopodia >4 μm in length shifting from 45% on glass to 13% on 2 kPa gels. Together, these results indicate that SSMs are mechanosensitive cells that adapt their morphology to changes in ECM rigidity.

### SSM mechanosensing of ECM rigidity alters the mobility of immune complexes

SSM mechanosensing of ECM rigidity also affected the formation of membrane ruffles on the dorsal cell surface. The images in Fig. 6A show that SSM dorsal membranes were replete with actin-rich ruffles when cells were cultured on collagen I-coated glass, and devoid of them when cells were cultured on collagen I-coated 2 kPa gels. Analysis of membrane ruffle formation showed that the fraction of SSMs with at least one ruffle decreased with ECM rigidity, from 100% of cells on glass to only 31% of cells on 2 kPa gels (Fig. 6B). This change in membrane topography did not impact the ability of SSMs to capture immune complexes, as the surface density (0.12 complexes/μm^2^) (Fig. 6C) and size (fluorescence intensity) (Fig. 6D) of immune complexes were unchanged. The loss of membrane ruffles and retention of immune complex density resulted in a loss of spatial association between immune complexes and ruffles on the dorsal surface (Fig. 6E). Because we previously found that the actin cytoskeleton constrains the lateral motion of immune complexes (Fig. 3 and 4), we hypothesised that ECM stiffness-induced alterations in membrane ruffle formation would also impact immune complex mobility. We therefore tracked immune complexes on the surfaces of live SSMs cultured on collagen I-coated substrates of different stiffness and analysed the trajectories using SMAUG. The weighted mean diffusion constant of immune complexes was 56% lower when SSMs were cultured on 2 kPa gels (0.0057 μm^2^/s) compared to glass (0.0129 μm^2^/s) (Fig. 6F). Detailed analysis of the different mobility states showed that the fraction of immobile complexes (D_immobile_<0.001 μm^2^/s) increased from 0.4 on glass to 0.6 on 2 kPa gels, and the fraction of complexes with D≥0.03 μm^2^/s decreased from 0.31 on glass to 0.06 on 2 kPa gels. Thus, membrane ruffling and stiff ECM increases the lateral mobilty of immune complexes presented by SSMs.

**Fig. 6:**
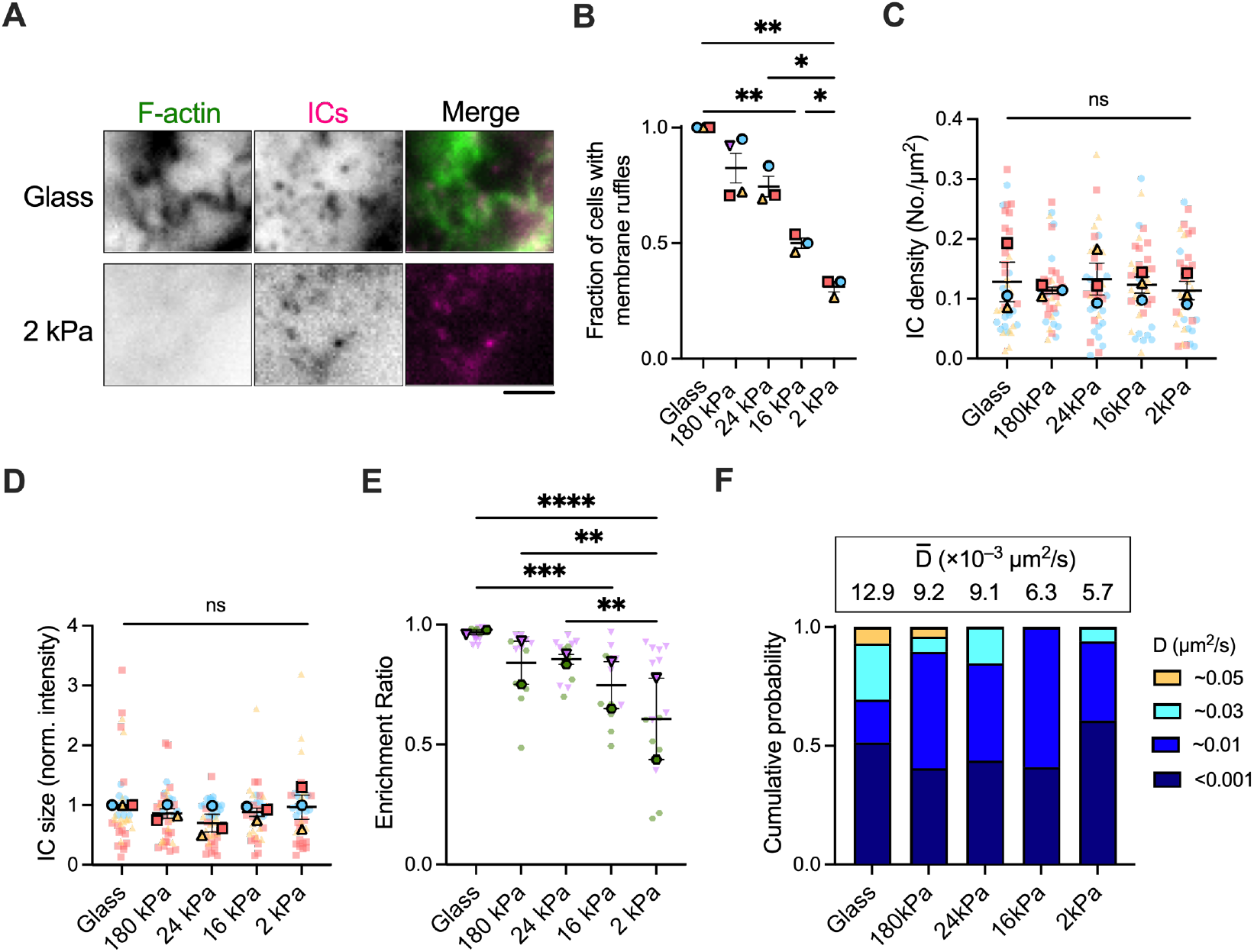
ECM rigidity alters membrane topography and immune complex mobility, but not the density or clustering of immune complexes. (A) Maximum intensity projection of three z-planes, totalling 0.6 μm depth, at the dorsal membrane of SSMs cultured on collagen I-coated glass (top row) or 2 kPa polyacrylamide gel substrates (bottom row). Cy3B-labelled immune complexes (magenta) are present on both cell membranes, but actin-rich membrane ruffles (stained with phalloidin-AF488, green) are visible only in the cell cultured on glass. Scale bar: 2 μm. (B) The fraction of cells on each substrate that have formed at least one membrane ruffle on the dorsal surface. N = 3 or 4 independent experiments. (C, D) Quantitation of the (C) surface density and (D) size of immune complexes presented by SSMs cultured on collagen I-coated glass or polyacrylamide gels of different stiffness. Each plain dot represents the mean value for all immune complexes on one cell and each outlined dot represents the mean value for all cells in one independent experiment (n = 8 to 19 cells per condition per experiment, N = 3 independent experiments). Data are colour-coded by experiment. Bars represent mean ± SEM. (E) Enrichment of immune complexes on actin-rich membrane ruffles on SSMs cultured on collagen I-coated gels of different stiffness (n = 5 to 14 cells per condition per experiment, N = 2 independent experiments). (F) Bar graphs showing the mean and weight fraction of each identified mobility state for immune complexes diffusing on cells cultured on collagen I-coated gels of different stiffness. Data are pooled from 2 independent experiments (2 kPa: 3095 trajectories from 7 cells, 16 kPa: 3020 trajectories from 4 cells, 24 kPa: 3100 trajectories from 5 cells, 180 kPa: 5340 trajectories from 8 cells, glass: 9160 trajectories from 7 cells.) The weighted mean of the diffusion constant for each condition, 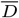, is given atop each bar. *p<0.05, **p<0.01, ***p<0.001, ****p<0.0001. ns, not significant.

## Discussion

Our study reveals a dynamic relationship between SSM membrane topography and immune complex presentation that is influenced by ECM mechanics. We show, using high-resolution fluorescence microscopy, that SSMs cultured on stiff collagen I-coated surfaces form prominent F-actin-based membrane protrusions. These protrusions enrich Fc*γ*R-presented immune complexes and promote their rapid movement on the cell surface. Culturing SSMs on soft collagen I-coated substrates depletes them of F-actin protrusions, leading to slower immune complex mobility. These findings demonstrate that physical aspects of immune complex presentation—namely, membrane topography and immune complex mobility—are not fixed features of SSMs, but rather dynamic qualities that change in response to mechanical forces transmitted from the ECM.

Our paper focuses on the role of SSMs as antigen-presenting cells, and as such, we should consider how SSMs interact with B cells. In the lymph node, B cells migrate to the sub-capsular sinus region to sample SSM surfaces for antigens (4). Most B cells spend <5 minutes in contact SSMs (2), so they must be efficient in their search. B cells rapidly scan their B cell receptors (BCRs) over antigen-presenting surfaces using actin-based membrane protrusions (31–33), which is a general mechanism employed by lymphocytes to locate antigens (34). We show here that SSMs, too, employ active surface topography to move immune complexes on their surfaces, suggesting that SSMs may survey B cell surfaces in return to facilitate encounters between immune complexes and BCRs.

Quantitative imaging of molecular interactions between B cells and SSMs *in vivo* is not currently possible. Instead, to dissect the molecular-scale events that underpin B cell activation, researchers have stimulated B cells with artificial surfaces to mimic cell-cell interactions in *vitro*. These experiments have revealed that, upon binding cognate antigen, B cells rapidly remodel their actin cytoskeleton to reposition BCRs and integrins to optimise signalling (35), and exert mechanical forces on BCRs to extract and internalise antigens (36). The ability of B cells to carry out these functions depends on physical properties of the antigen-presenting surface (37). For example, *in vitro* experiments have shown that B cells interacting with antigens on fluid planar lipid bilayers (D_Ag_ ≈ 4 μm^2^/s) (38) easily assemble large BCR-antigen microclusters that signal robustly (39). In contrast, B cells encountering the same antigens tethered to glass coverslips (D_Ag_ = 0 μm^2^/s) form much smaller clusters and have an attenuated response. In this paper, we reveal that neither of these scenarios is an accurate representation of antigen presentation by SSMs. Using single particle tracking and quantitative analysis of the trajectories, we show that immune complexes presented by SSMs frequently transition between immobile (D<0.001 μm^2^/s) and moderate-mobility (0.03-0.05 μm^2^/s) states in a way that depends upon actin. These diffusion constants are close to those of BCRs (D_IgM-BCR_ = 0.03 μm^2^/s and D_IgD-BCR_ = 0.003 μm^2^/s) (40), suggesting that SSMs may resist the lateral transport of BCR-antigen complexes to inhibit microcluster growth. Speculatively, certain immune complex mobility states may also be more favourable for BCR microcluster formation. Moreover, we show that SSMs present (multivalent) immune complexes at low average densities (0.12 complexes/μm^2^), in contrast to the very high antigen densities commonly used *in vitro* (50 to 4000 molecules/μm^2^) (13, 35, 41). We suspect that presenting multivalent antigens at low density and with low mobility will maintain low-avidity BCR-antigen interactions to promote stringent B cell discrimination of antigen affinities (42). These findings invite the development of artificial surfaces that better imitate physical characteristics of real antigenpresenting cells for future *in vitro* studies of B cell activation.

An important finding in our experiments is the discovery that SSMs are mechanosensitive cells that respond to ECM rigidity differences by altering F-actin architecture to modify cell shape, membrane topography, and immune complex mobility. The *in vivo* relevance of SSM mechanotransduction stems from the fact that in response to immunological challenge, lymph nodes rapidly and massively increase their volume, which requires remodelling of stromal and ECM networks. The network tension remains the same or even decreases during early stages of lymph node swelling (43), but increases as the lymph node continues to expand (44, 45). We speculate that SSMs in contact with ECM will sense these mechanical changes and alter immune complex presentation. Our experiments suggest that increasing ECM tension may induce the formation of SSM F-actin structures, including dorsal membrane ruffles that enhance immune complex mobility and filopodia that extend into the B cell follicle to deliver antigens (2, 4–6). We do not yet know the mechanisms underpinning SSM sensing of ECM mechanics, but it is conceivable that integrin-mediated mechanotransduction activates small GTPase activity to alter filopodia formation and membrane ruffling (46–48). Another point to consider in future studies will be the contributions of different ECM ligands to SSM activation, as different integrin types exhibit different bond dynamics and consequently responses to ECM rigidity (49). Our findings overall suggest that changes to ECM tension in lymph nodes may provide a mechanical stimulus to SSMs that alters physical aspects of antigen presentation to B cells, thereby shaping B cell activation responses. In summary, we have demonstrated that SSMs can be enriched from lymph nodes with IgG antibody complexes, and identified in *in vitro* cultures with high-throughput, multi-colour fluorescence imaging. SSMs have distinct morphological and topographical features that are linked to immune complex presentation and alter dynamically in response to ECM rigidity. Understanding the mechanisms that regulate SSM mechanosensing and the spatial organisation of immune complexes on the cell surface, especially *in vivo*, will be needed to determine their exact roles in supporting B cell responses. Immunisation strategies that consider ECM tension in addition to antigen accumulation in lymph nodes may provide new routes to elicit antibodies with desired breadth and specificity (50, 51), for instance by targeting stromal cell contractility to alter ECM tension (52).

## Materials and Methods

### Mice

C57BL/6 mice were purchased from Charles River and housed in the King’s College London Biological Services Unit under specific-pathogen-free conditions. All mice were 6-16 weeks of age, and both males and females were used. Mice were maintained and treated following guidelines set by the UK Home Office and the King’s College London Ethical Review Panel.

### SSM isolation and culture for imaging experiments

Lymph nodes (superficial and deep cervical, brachial, axillary, mesenteric, and inguinal) were harvested from five C57BL/6 mice and teased apart using 25G needles in a 35 mm Petri dish containing 1 mL DMEM medium. Another 1 ml DMEM medium containing 0.625 mg/ml DNase1 (Roche) and 0.26 U Liberase DH (Sigma) was added to the cells and incubated for 45 minutes at 37 °C. Released cells were collected, and the tissue was digested a second time with fresh reagents. The cells were pooled and passed through a 70 μm cell strainer and pelleted by centrifugation for 7 minutes at 300×g. The cell pellet was resuspended in 200 μl ice-cold DMEM and incubated sequentially on ice with purified rat anti-mouse FDC-M1 (BD Biosciences) or rat IgG2c, *κ* isotype control antibody (1.6 μg antibody per 2 × 10^7^ cells) for 1 hour, 1 μg biotinylated mouse anti-rat Ig, *κ* light chain (BD Biosciences) antibody for 40 minutes, and 50 μl anti-biotin microbeads (Miltenyi Biotec) for 20 minutes. The cells were pelleted (5 minutes at 300×g) and resuspended in full DMEM (DMEM supplemented with 10% heat-inactivated fetal calf serum, 20 mM HEPES, 0.2 mM MEM non-essential amino acids, and penicillin-streptomycin-glutamine, all from Gibco) between each incubation. Following a final wash and resuspension of the cell pellet in MACS buffer (PBS, pH 7.4, 1 mM EDTA, 5% BSA), cells were isolated by positive selection using an LS Column and MidiMACS Separator (Miltenyi Biotec). Cells were resuspended in full DMEM and plated onto collagen I-coated glass or polyacrylamide gels in FluoroDish cell culture dishes (10-mm well; World Precision Instruments) at a density of 3 × 10^5^ cells per dish. Cells were cultured at 37 °C with 5% CO_2_. The media was exchanged at 48 hours to remove dead lymphocytes and cell debris. Cells were imaged on days 5-7.

For actin perturbation experiments, cells were exchanged into warm Hank’s balanced salt solution (+Ca^2+^, +Mg^2+^; Gibco) supplemented with 0.01% BSA (HBSS 0.01% BSA) and mycalolide B, jasplakinolide, CK-666, SMIFH2, Y-27632, or blebbistatin at final concentrations listed in Supplementary Table 2. Control cells received an equivalent amount of DMSO diluted in warm HBSS 0.01% BSA.

### Immune complex generation

Blood was collected from a mouse by cardiac puncture and allowed to clot for 30 minutes at room temperature. The clot was removed by centrifuging at 2000×g for 10 minutes at 4 °C. The liquid component (serum) was divided into 10 μl aliquots on ice and stored at −20 °C. Immune complexes were generated by mixing 10 μl of the mouse serum, as a source of complement, with 0.5 μg Cy3B-labelled goat IgG1 anti-mouse Ig*κ* (Southern Biotech; labelled in-house), 0.375 μg donkey anti-goat IgG (H+L) (Jackson ImmunoResearch), and 40 μl of GVB++ buffer (Complement Technology) for 30 minutes at 37 °C.

### Antibodies for imaging and flow cytometry

Antibodies for cell enrichment, flow cytometry, and fluorescence imaging are listed in Supplementary Table 1. In-house labelling of antibodies was achieved by mixing NHS ester-coupled dyes with the antibody at a 4:1 dye:antibody ratio in sodium bicarbonate buffer, pH 8.2, for 30 minutes at room temperature and then removing unbound dye molecules using 7 kDa MWCO Zeba desalting columns (Pierce).

### Cell characterisation by flow cytometry

Single cells from the whole lymph node and positive selected fraction were split into 10^6^ cells/tube and blocked with anti-mouse CD16/32 in PBS for 20 minutes at 4 °C and surface stained for 30 minutes at 4 °C with labelled primary antibodies or their isotype controls (see Supplementary Table 1). Propidium iodide was used to exclude dead cells. Flow cytometry was performed on a BD FACS Canto II. Data were analysed using FlowJo software (BD Biosciences).

### Imaging and image processing

Widefield z-stack and timelapse fluorescence images were acquired using a Nikon TiE TIRF microscope equipped with a 100×, 1.49-NA oil-immersion objective (Nikon), a motorised stage with an integrated piezo Z-drive (MS-2000; Applied Scientific Instrumentation), and active Z-drift correction (Perfect Focus System; Nikon). The microscope was controlled by a high-speed TTL, I/O, DAC controller (Triggerscope 4; Cairn Research) integrated into MicroManager Software. Illumination was supplied by a MultiLine LaserBank (Cairn Research) fitted with 405-, 488-, 561-, and 640-nm diode lasers (Coherent OBIS). The beams were aligned into a single-mode fibre and coupled to an iLas2 Targeted Laser Illuminator (Gataca Systems), which produces a 360° spinning beam with an adjustable illumination angle. Laser beams were passed through a laser quadband (405/488/561/640nm) filter set for TIRF applications (TRF89901v2-ET; Chroma) before illuminating the sample. Emitted fluorophores were filtered by appropriate single-band emission filters (Chroma) using a filter wheel (OptoSpin; Cairn Research) and then captured onto a back-illuminated sCMOS camera (Prime 95B sCMOS; Teledyne Photometrics). For live-cell imaging, a constant temperature of 37 °C was maintained using a cage incubator (Oko-lab). Images were processed and analysed using Fiji (53) and Icy v1.9.9.1 (54). Briefly, images from each channel were aligned and cropped to remove poorly illuminated areas at the edges. Images were then background-subtracted and flatfield corrected before analysis.

### Plots and statistics

Data for individual cells and the mean values per experiment were plotted together as SuperPlots (55). Statistical differences were determined using one-way ANOVA with Tukey’s multiple comparisons test. Statistical tests were performed using GraphPad Prism software (version 9), and differences were considered to be statistically significant at p ≤ 0.05.

### Preparation of glass substrates

Stock solutions of collagen were prepared by dissolving 10 mg collagen I (rat tail-derived; Roche) in 3.3 mL of 0.2% (v/v) acetic acid in water. The resulting 3 mg/ml stock solution was stored at 4 °C. On the day of an imaging experiment, the collagen was diluted to 30 μg/ml in PBS, pH 7.4, and incubated on the 10-mm FluoroDish glass surface for 3 hours at 37 °C. The collagen was aspirated, and the surfaces were incubated with full DMEM for 20 to 30 minutes at 37 °C, before cells were seeded.

### Preparation of polyacrylamide gels

The glass surfaces of 10-mm FluoroDishes were each incubated with 100 μl of 0.1 M sodium hydroxide for 5 minutes, washed with ultrapure water (Arium), and dried. The surfaces were then incubated with 100 μl of 1.5% (v/v) (3-aminopropyl)trimethoxysilane in ultrapure water for 30 minutes at room temperature. The surfaces were washed again and incubated with 100 μl of 0.5% glutaraldehyde in PBS for 30 minutes, washed with ultrapure water, and dried. To generate gels of different stiffness, different concentrations of acrylamide and bisacrylamide (see Supplementary Table 3) were mixed with 0.1% ammonium persulfate and 0.1% TEMED (N,N,N’,N’-tetramethylethylenediamine) in degassed 10 mM HEPES, pH 7.0. Two microlitres of the solution were promptly placed in the centre of the FluoroDish glass surface and covered with a 9-mm round glass coverslip (made hydrophobic by prior treatment with Rain-X). Two magnets were used to hold the glass-gel-glass sandwich together during polymerisation to ensure uniform gel thickness. After 30 minutes the polymerisation was complete, the round coverslips were removed, and the gels were soaked in 50 mM HEPES, pH 7.0. To coat the gel surfaces with collagen, they were first incubated with 100 μl of 0.5 mg/ml sulfo-SANPAH (sulfosuccinimidyl 6-(4’-azido-2’-nitrophenylamino)hexanoate); Thermo Scientific) in 10 mM HEPES, pH 8.5, and exposed to 6 W of 365 nm UV light (UVP UVL-56 handheld UV lamp) until the solution turned from orange to brown. Excess sulfo-SANPAH was removed by washing with 50 mM HEPES, pH 8.5. Crosslinked gels were then incubated with 100 μl of 30 μg/ml collagen I diluted in 50 mM HEPES, pH 8.5, for 2 hours at 37 °C. Gels were washed with full DMEM and incubated for 20-30 minutes at 37 °C before adding cells.

### Measurements of polyacrylamide gel stiffness

The stiffness (Young’s modulus, E) of polyacrylamide gels was measured by nanoindentation using an atomic force microscopy (BioScope Resolve, Bruker). Briefly, cantilevers with pyramidal silicon nitride tips with an effective half-angle *ϑ* of 18°, nominal spring constant of k=0.03 N m^-1^, nominal tip radius of 20 nm, and minimal tip height of 2.5 μm (MLCT-D, Bruker) were used. The AFM was calibrated using the deflection sensitivity and actual spring constant of the cantilevers. To determine the deflection sensitivity of the cantilevers, a deflection/force curve for the approach on a glass surface was recorded. The actual spring constant was found through thermal tuning using the simple harmonic oscillator model (56). Two gels per stiffness were measured. For each gel, measurements were obtained from four areas around the centre, spaced at least 4 μm apart. Per area, 64 (8×8) force-displacement curves with a ramp size of 5 μm and ramp speed of 10 μm s^-1^ were recorded. The Young’s modulus was then computed using a model for force-indentation relationships of a four-sided pyramidal tip (57). All the force curves were processed in this way with a custom-written MATLAB code.

### Immunofluorescence imaging of fixed cells

For immunofluorescence imaging, cells were exchanged into icecold HBSS 0.1% BSA, incubated with immune complexes for 10 minutes on ice and fixed with 4% paraformaldehyde. Cells were then blocked with anti-mouse CD16/32 and stained for surface markers using antibodies at a final concentration of 1 μg/ml for 30 minutes at room temperature. For staining cytoplasmic molecules, cells were permeabilised with 0.1% Triton X-100 (Alfa Aesar) for 5 minutes in HBSS 0.1% BSA, blocked with 5% (v/v) normal mouse serum (Jackson ImmunoResearch), and incubated with the antibody for intracellular staining. F-actin was stained using Alexa Fluor 488 phalloidin (Invitrogen). Cells were fixed again before imaging.

Because jasplakinolide competitively inhibits phalloidin binding to F-actin, preventing the combined use of these drugs during imaging experiments (58), we used a silicon rhodamine (SiR)-jasplakinolide conjugate. At low concentrations SiR-jasplakinolide can be used as a cell-permeable probe to study actin dynamics, but at the 100 nM concentration we used, it potently induced actin polymerisation and inhibited actin turnover while also allowing us to visualise actin filaments.

### Analysis of cell morphology

The cell body was segmented based on the intensity of the F-actin labelling. The cell shape, circularity and volume were determined in Icy using the ROI statistics plugin. The podosome size and filopodia length were measured in Fiji. Briefly, F-actin signals corresponding to podosomes were thresholded, binarized, and applied as a mask to the F-actin raw image, and the Analyse Particles plugin was used to quantify podosome number, size, and fluorescence intensity. Filopodia lengths in SSMs cultured on different substrates were measured manually, using the Fiji line selection tool, by a researcher knowledgeable of the actin structures but blinded to the data set (i.e., which substrate cells were adhered to).

### Fluorescence intensity-based colocalisation analysis of immune complexes and cytoskeletal filaments

Our approach to quantify the association of immune complexes (spots) and F-actin (filaments) was inspired by Sun et al. (59). Briefly, two-colour z-stack epifluorescence images (phalloidin and immune complex channels) were processed and analysed by a user-guided pipeline implemented in Fiji (53). There were three steps to the analysis: (1) identify the z-positions of the dorsal membrane ruffles, (2) remove low-frequency signals from F-actin and immune complex channels, and (3) quantify the spatial association of immune complexes and F-actin.

Because dorsal membrane ruffles and immune complexes in different regions of the cell come into focus at different z-stack positions, six regions of interest (ROIs) were analysed for each cell. The Sobel transform and Canny-Deriche operator were applied to the phalloidin channel, and the three adjacent z-slices with the highest integrated values (sharpest frames), corresponding to the z-slices with dorsal membrane ruffles, were identified. A maximum intensity projection of the three z-slices from the phalloidin raw image stack was generated and used for further analysis. A maximum intensity projection of the same three z-slices was produced from the immune complex raw image stack.

Next, segmented images of dorsal membrane ruffles and immune complexes were generated. For each channel (phalloidin and immune complexes), low-frequency signals were subtracted (using a Gaussian bandpass filter in the phalloidin channel and a Laplacian of Gaussians filter in the immune complex channel) and then each channel was thresholded, binarised, and subsequently applied as a mask to the raw phalloidin and immune complex channels to exclude background fluorescence signal. The immune complexes were identified, and the raw integrated density of each complex quantified, using the Analyse Particles plugin. The enrichment ratio of immune complexes on actin filaments, R, was then calculated for each ROI as the summed intensity of complexes that are “on actin”, I_c,on_, divided by the summed intensity of all immune complexes, I_c,on+off_, or

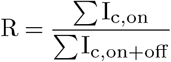

The same procedure was used to calculate the enrichment ratio for immune complexes with microtubules, vimentin, and nestin. For these experiments, antibodies targeting the cytoskeletal proteins were used in place of phalloidin.

### Live-cell imaging and analysis

Cells cultured on collagen I-coated substrates were incubated with fluorescent immune complexes for 10 minutes on ice, exchanged into warm HBSS 0.1% BSA, and imaged at 37 °C. Streamed timelapse images were acquired at a frame rate of 20 Hz for 15 seconds at a single plane focused on the dorsal cell membrane. Images were background-subtracted and corrected for photobleaching using either the histogram matching method (immune complex channel) or the exponential fitting method (Lifeact-GFP channel) in Fiji (60). A Difference of Gaussians filter was applied to the immune complex image (σ_1_: 3.00 and σ_2_: 1.50). The particles were detected and tracked with sub-pixel localisation using the LAP tracker in the Trackmate plugin for Fiji (61), with a maximum linking distance of 500 nm, a maximum gap-closing distance of 250 nm, and a maximum gap of 2 frames. For the kymograph analysis, single-particle signals were extracted along the immune complex trajectory. Kymographs were then generated from videos of 300 frames using the Multi Kymograph plugin for Fiji, with a line width of 1 pixel. The evolution of fluorescence intensity along the immune complex trajectories was plotted versus time to generate the kymographs for both the immune complex and F-actin channels. The MSD of each immune complex diffusing in two-dimensions, in the plane of the plasma membrane, was computed according to the formula (62)

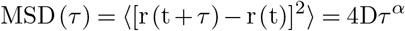

where *τ* = 50 ms is the frame time, [r(t + *τ*) — r (t)] is the immune complex displacement during time interval *τ*, D is the apparent diffusion constant, and *α* is the anomalous scaling exponent, with *α* =1 characterised as Brownian (normal) diffusion, *α* < 1 as subdiffusion, and *α* > 1 as superdiffusion. Trajectories of 10 spots or longer were used for the analysis. Linear trajectories due to active transport of internalised immune complexes were excluded from the analysis.

## ACKNOWLEDGEMENTS

We thank Stephen Tovey (Cairn Research) for expert assistance with assembling the microscope system and integrating it with MicroManager, Dessi Malinova for providing mouse serum, and Felix Gehres and Sergi Garcia-Manyes for assistance with measuring polyacrylamide gel stiffness. We thank the KCL Biological Services Unit and flow cytometry platform. This work was supported by a BBSRC research grant (BB/S007814/1), a BBSRC sLoLa (BB/V003518/1), and a Royal Society research grant (RGS\R2\180333) to KMS.

## AUTHOR CONTRIBUTIONS

MI designed and performed the experiments, analysed the data, and helped prepare the manuscript. ATB performed flow cytometry experiments, analysed the data, and provided advice. CC performed flow cytometry experiments and analysed the data. HCWM developed image analysis pipelines. FA analysed imaging data. RK analysed flow cytometry data and provided LifeAct-GFP mice. APC designed the experiments and provided reagents and advance. KMS designed the experiments, supervised the research, and wrote the manuscript. All authors contributed to revising and editing the manuscript.

---

**Fig. S1:**
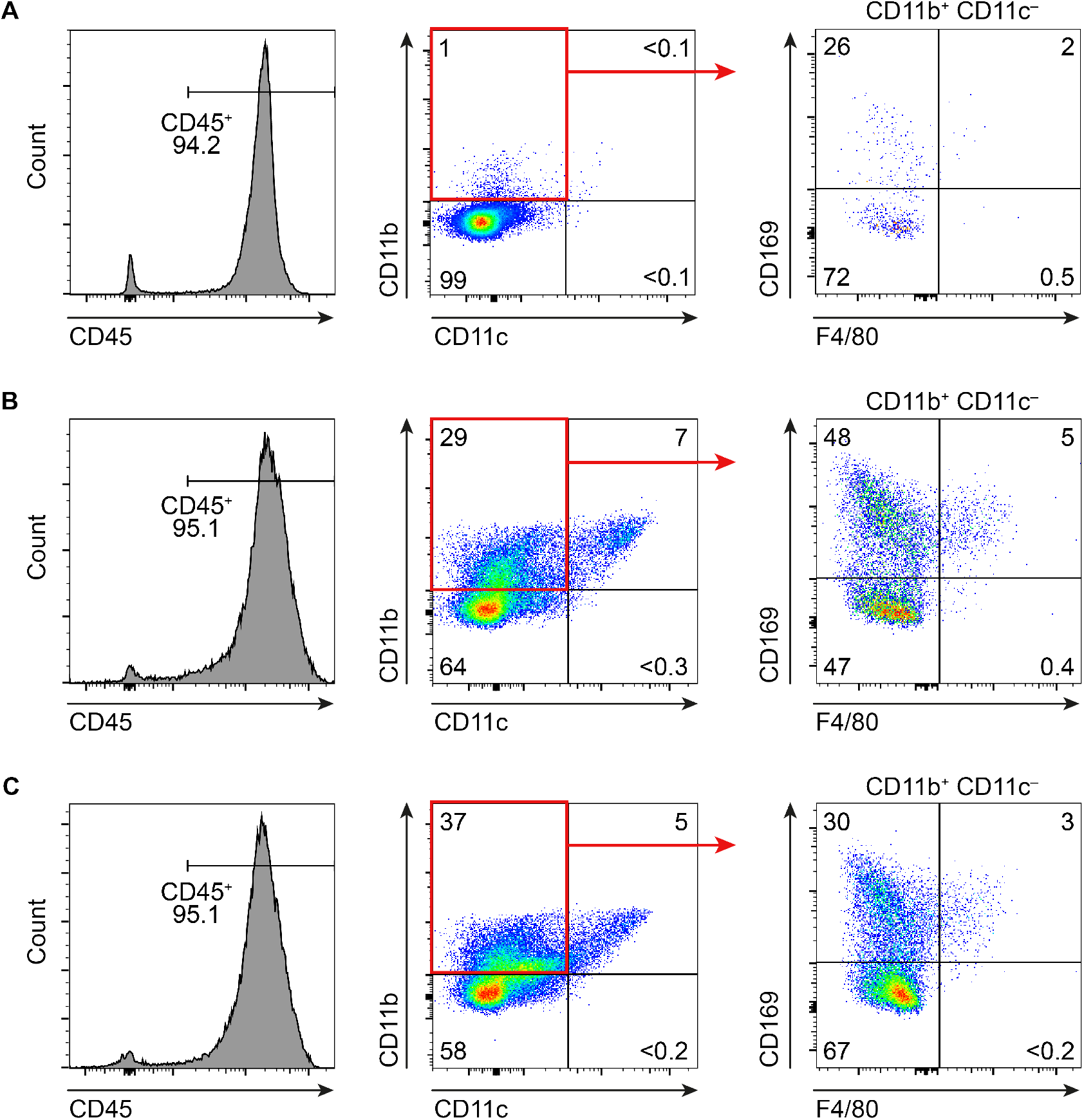
Isolation and identification of SSMs. Pseudocolour flow cytometry plots of live single-cell suspensions from (A) whole lymph nodes, and (B,C) enzyme-digested lymph nodes positively enriched by (B) an FDC-specific antibody (rat IgG2c, *κ* anti-mouse FDC-M1) or (C) a rat IgG2c, *κ* isotype control antibody, both complexed with a biotinylated mouse IgG2a anti-rat Ig *κ* secondary antibody and captured by anti-biotin microbeads. Cells were stained with monoclonal antibodies to CD45, CD11b, CD11c, CD169, and F4/80 (see Supplementary Table 1). The proportions of all cell populations are indicated on the plots. Data are representative of three experiments.

**Fig. S2:**
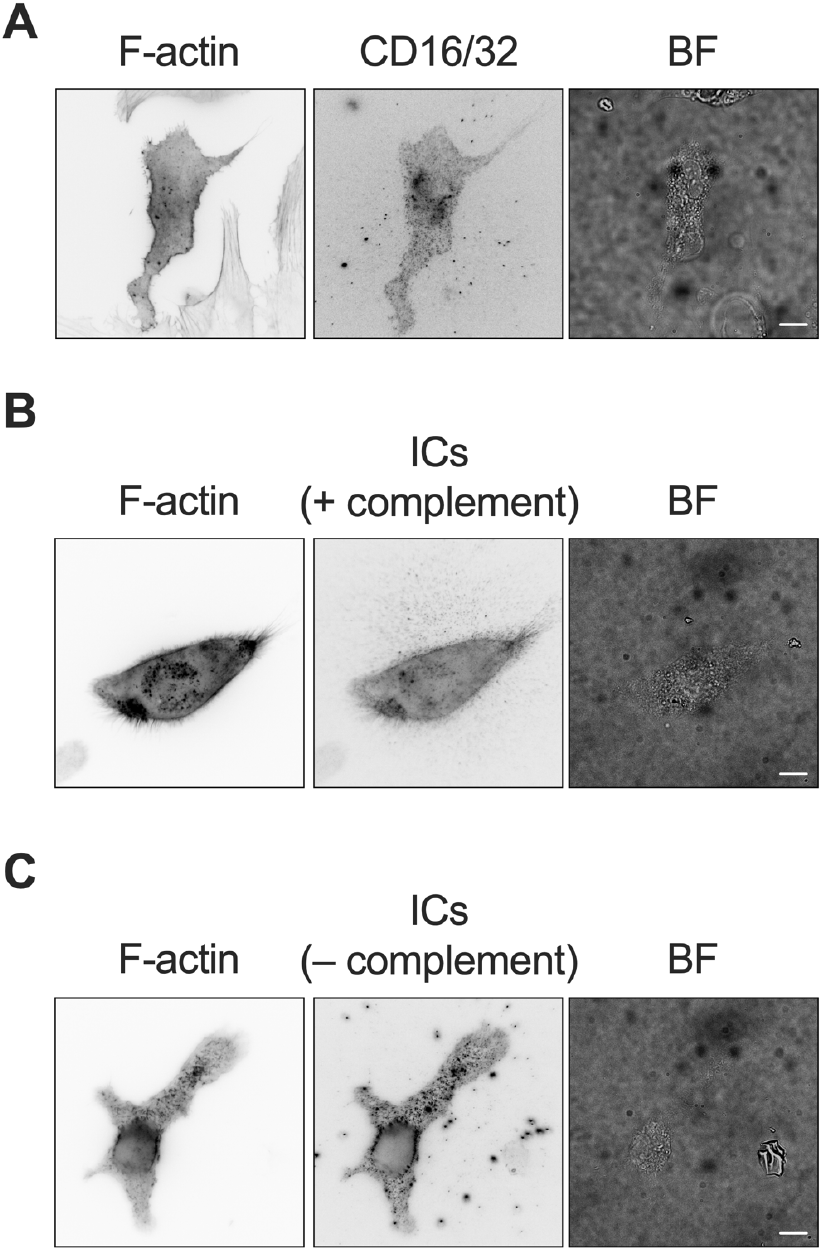
SSMs use Fc*γ*Rs to present immune complexes. (A) Immunofluorescence staining with anti-CD16/32 (Fc*γ*RIII/Fc*γ*RII). Both SSMs (F-actin^+^ CD16/32^+^) and non-SSMs (F-actin^+^ CD16/32^-^) are visible in this image. (B, C) The ability of SSMs to capture and present immune complexes does not depend upon complement. IgG antibody complexes are captured equally well when (B) they are (+) and (C) are not (-) fixed with complement. Scale bars: 10 μm.

**Fig. S3:**
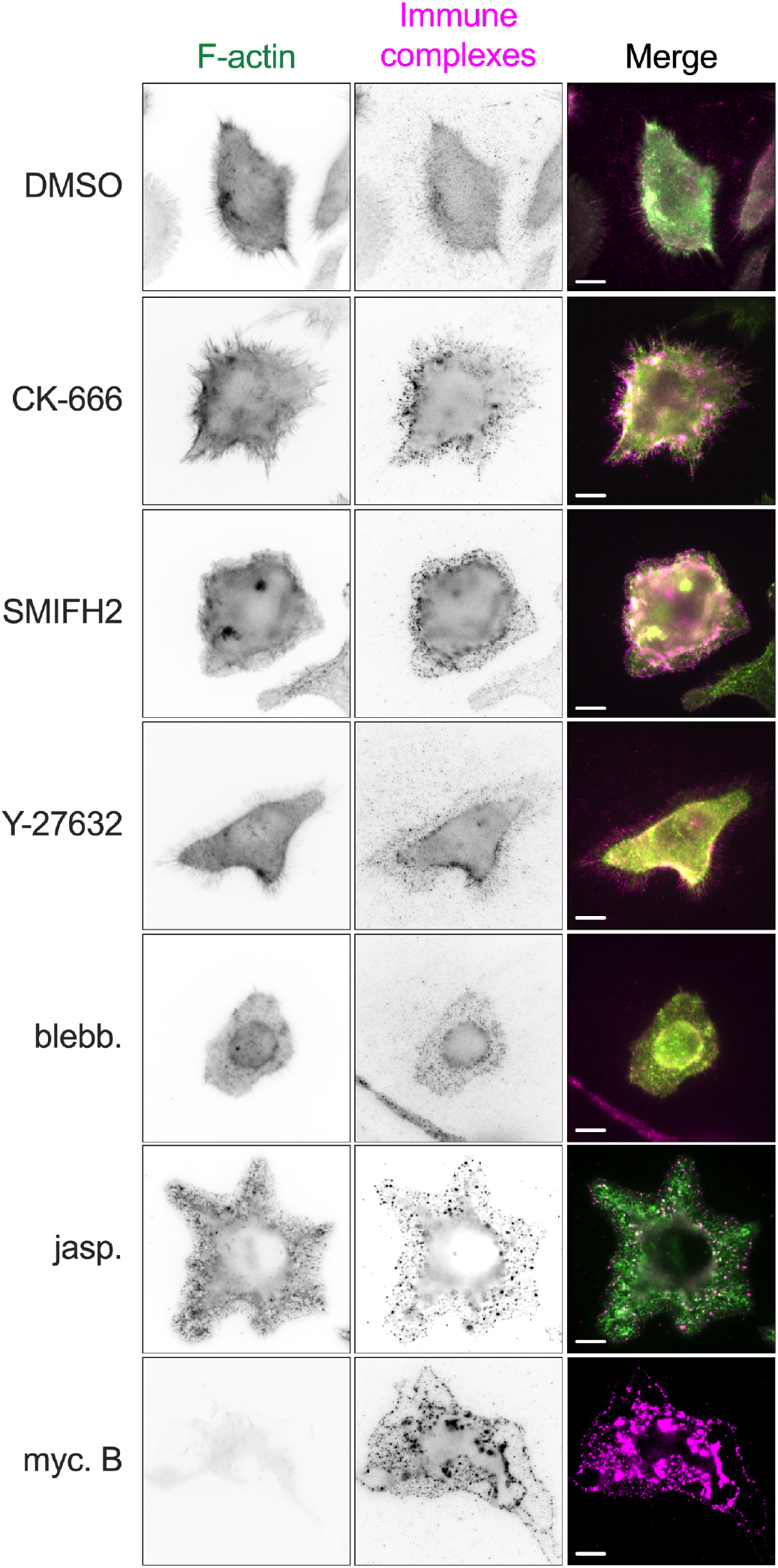
Inhibition of SSM actin dynamics and contractility. F-actin (phalloidin-AF488) and immune complex (Cy3B-labelled) staining of SSMs treated with DMSO or inhibitors of the following targets: Arp2/3 (CK-666), formins (SMIFH2), Rho-associated kinase (Y-27632), myosin II (blebbistatin), actin disassembly (fluorogenic form of jasplakinolide, SiR-Actin), and actin assembly (mycalolide B). Inhibitor concentrations are listed in Supplementary Table 2. Scale bars: 10 μm.

**Fig. S4:**
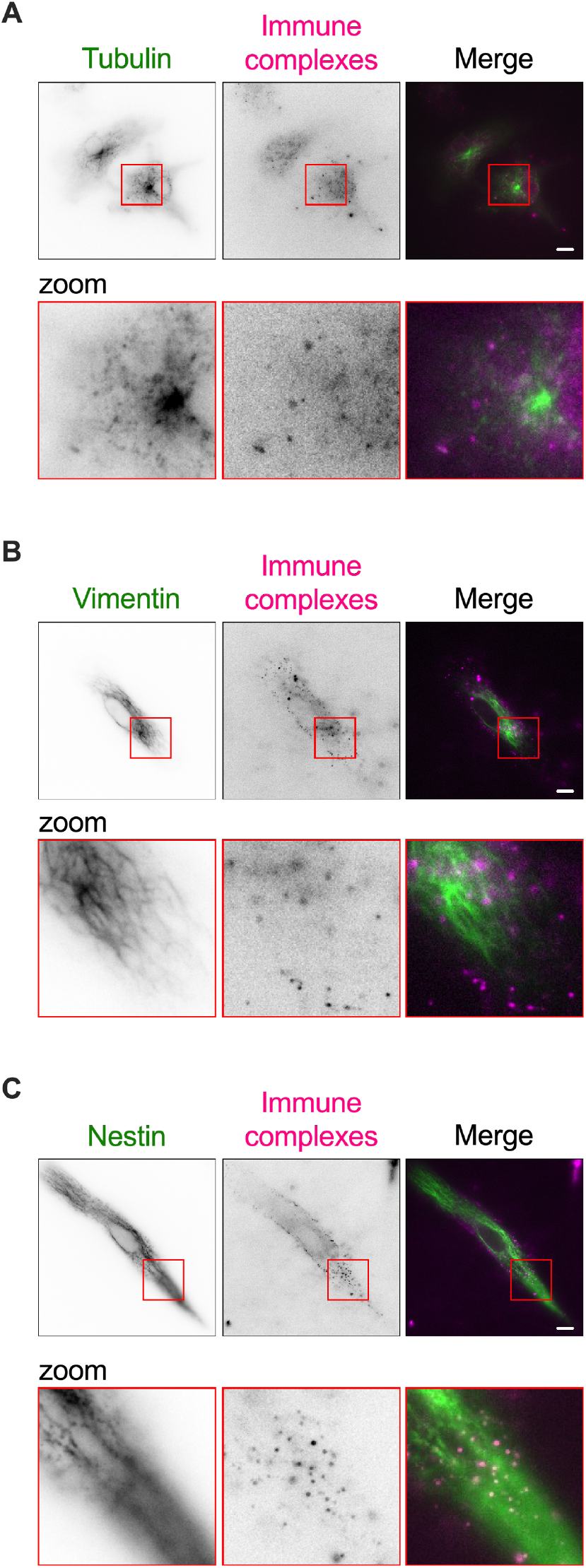
Immune complexes presented by SSMs do not associate with microtubules or intermediate filaments. SSMs were cultured on collagen I-coated glass, labelled with Cy3B-immune complexes, and stained for (A) tubulin (z-position: 2 μm), (B) vimentin (z-position: 2.4 μm), and (C) nestin (z-position: 3.6 μm). Scale bars: 10 μm.

**Fig. S5:**
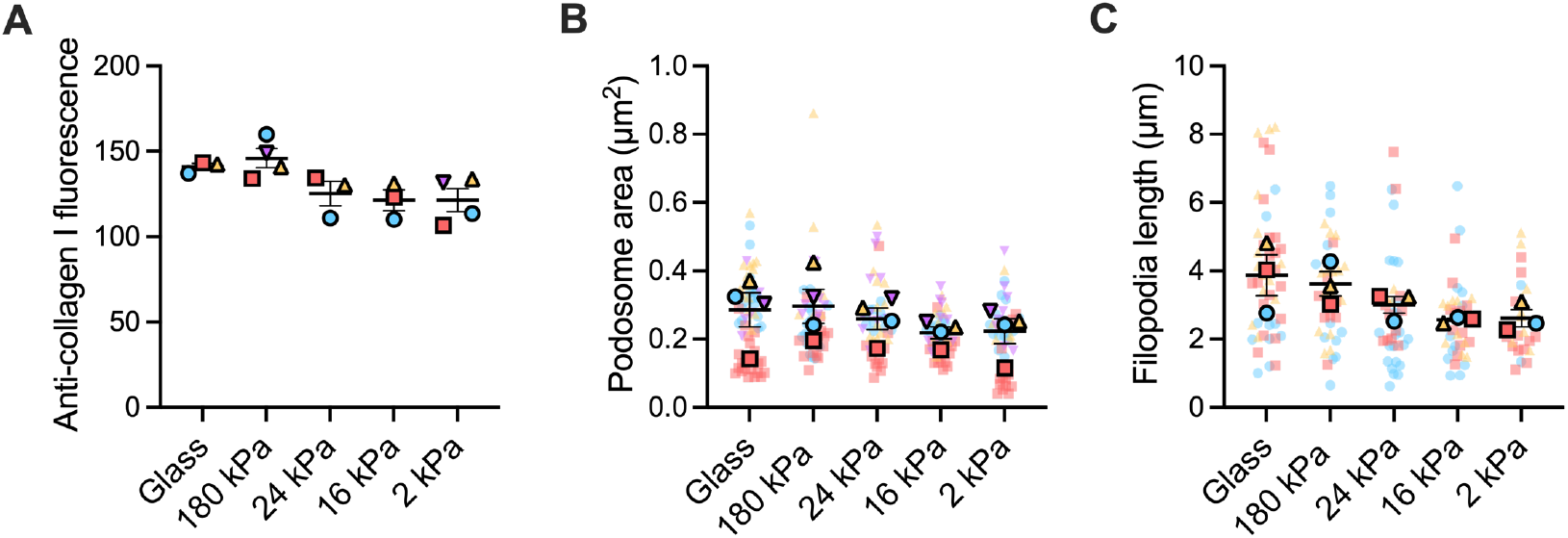
Characterising polyacrylamide gels and cell morphology. (A) Collagen I-coated glass and gel substrates were stained with a fluorescent anti-collagen I antibody and imaged with epifluorescence microscopy. The fluorescence intensity was the same across gels of different rigidity, indicating a consistent surface density of collagen I. Data are the mean fluorescence intensities from 3 or 4 gels of each stiffness. (B) Podosome areas and (C) filopodia lengths for SSMs cultured on collagen I-coated substrates of different stiffness. Each plain dot represents one cell and each outlined dot represents the mean value for one independent experiment (n = 2 to 23 cells per condition per experiment, N = 3 or 4 independent experiments. Bars represent mean ± SEM.

**Supplementary Table 1:** Antibodies used in this study

| Antibody | Clone | Final conc. | Supplier | Cat. No. | Assay |
| --- | --- | --- | --- | --- | --- |
| CD45-AF647 | 30-F11 | 0.05 µg/ml | Biolegend | 103123 | Flow cytometry/Imaging |
| CD169-FITC | 3D6.112 | 0.05 µg/ml | Biolegend | 142405 | Flow cytometry/Imaging |
| CD11b-PE/Cy7 | M1/70 | 0.04 µg/ml | eBioscience | 25-01112-81 | Flow cytometry |
| CD11c-APC/Cy7 | N418 | 0.04 µg/ml | Biolegend | 117323 | Flow cytometry |
| F4/80-Pacific Blue | BM8 | 0.05 µg/ml | Biolegend | 123123 | Flow cytometry/Imaging |
| CD45-APC/Cy7 | 30-F11 | 0.05 µg/ml | Biolegend | 103115 | Flow cytometry |
| Rat IgG2a-FITC iso ctrl | eBR2a | 0.05 µg/ml | eBioscience | 11-4321-71 | Flow cytometry/Imaging |
| Rat IgG2a-Pacific Blue iso ctrl | RTK2758 | 0.05 µg/ml | Biolegend | 400527 | Flow cytometry/Imaging |
| Rat IgG2a-APC/Cy7 iso ctrl | RTK2758 | 0.04 µg/ml | Biolegend | 400523 | Flow cytometry |
| Rat IgG2b-AF647 iso ctrl | RTK4530 | 0.05 µg/ml | Biolegend | 400626 | Flow cytometry/Imaging |
| Rat IgG2b-PE/Cy7 iso ctrl | RTK4530 | 0.04 µg/ml | Biolegend | 400617 | Flow cytometry |
| Rat IgG2b-APC/Cy7 iso ctrl | RTK4530 | 0.05 µg/ml | Biolegend | 400623 | Flow cytometry |
| Arm. Hamster-APC/Cy7 iso ctrl | HTK888 | 0.04 µg/ml | Biolegend | 400927 | Flow cytometry |
| CD11b-biotin | M1/70 | 0.05 µg/ml | BD | 553309 | Imaging |
| CD16/32 (labelled in-house) | 2.4G2 | 0.05 µg/ml | BD | 553142 | Imaging |
| CD11c-BV605 | N418 | 0.05 µg/ml | Biolegend | 117333 | Imaging |
| CD68-AF488 | FA11 | 0.05 µg/ml | Biolegend | 130712 | Imaging |
| CD209b (labelled in-house) | eBio22D1 | 0.05 µg/ml | Invitrogen | 14-2093-81 | Imaging |
| Collagen I (labelled in-house) | COL-1 | 0.05 µg/µl | Invitrogen | PA-95137 | Imaging |
| FDC-M1 | FDC-M1 | see methods | BD | 551320 | Cell enrichment |
| Rat IgG2c, k iso ctrl | A23-1 | see methods | BD | 553982 | Cell enrichment |
| Biotin anti-Rat Ig, $\kappa$ | MRK-1 | see methods | BD | 553871 | Cell enrichment |
| Goat anti-mouse $\kappa$ IgG | polyclonal | see methods | SouthernBiotech | 1050-01 | Immune complex |
| Donkey anti-goat IgG (H+L) | polyclonal | see methods | Jackson | 705-005-147 | Immune complex |

**Supplementary Table 2:** Inhibitors used in this study

| Inhibitor | Target | Final conc. | Supplier | Cat. No. |
| --- | --- | --- | --- | --- |
| Mycalolide B | Actin severing | 3 µM | Santa Cruz | SC-358736 |
| Jasplakiolide (SiR-Actin) | Actin stabilisation | 100 nM | Spirochrome | SC001:SiR-actin kit |
| CK-666 | Arp2/3 | 100 µM | Sigma | SML0006 |
| SMIFH2 | Formin FH2 domain | 20 µM | Merck | 344092 |
| Y-27632 | Rho-associated protein kinase | 1 µM | Cambridge Bioscience | SM02-1 |
| (S)-nitro-blebbistatin | Non-muscle myosin II ATPase | 100 µM | Cambridge Bioscience | CAY24171 |

**Supplementary Table 3:** Polyacrylamide gel rigidities measured by atomic force microscopy (AFM)

| % Acrylamide | % Bis-Acrylamide | Young's modulus (kPa) (mean ± SEM) | No. gels measured |
| --- | --- | --- | --- |
| 4.96 | 0.05 | 1.88 ± 21.3 | 2 |
| 5.56 | 0.27 | 16.25 ± 0.77 | 2 |
| 7.06 | 0.20 | 24.24 ± 0.65 | 2 |
| 9.91 | 0.43 | 176.85 ± 0.21 | 2 |

**Movie S1.**

Raw epifluorescence timelapse of Cy3B-labelled immune complexes diffusing in the plasma membrane of a mouse subcapsular sinus macrophage expressing Lifeact-GFP. The still images in Figure 3, B and C, are taken from this movie. Image size: 252 x 69 pixels, pixel size: 110 nm.

**Movie S2.**

Raw epifluorescence timelapse of Cy3B-labelled immune complexes diffusing in the plasma membrane of a mouse subcapsular sinus macrophage treated with mycalolide B. Image size: 90 x 90 pixels, pixel size: 110 nm.

**Movie S3.**

Raw epifluorescence timelapse of Cy3B-labelled immune complexes diffusing in the plasma membrane of a mouse subcapsular sinus macrophage treated with fluorogenic jasplakinolide (SiR-Actin). Image size: 180 x 90 pixels, pixel size: 110 nm.

## Bibliography

1. Isabelle Mondor, Myriam Baratin, Marine Lagueyrie, Lisa Saro, Sandrine Henri, Rebecca Gentek, Delphine Suerinck, Wolfgang Kastenmuller, Jean X. Jiang, and Marc Bajénoff. Lymphatic Endothelial Cells Are Essential Components of the Subcapsular Sinus Macrophage Niche. Immunity, 50(6):1453–1466.e4, June 2019. ISSN 10747613. doi: 10.1016/j.immuni.2019.04.002.

2. Yolanda R Carrasco and Facundo D Batista. B cells acquire particulate antigen in a macrophage-rich area at the boundary between the follicle and the subcapsular sinus of the lymph node. Immunity, 27(1):160–171, July 2007. doi: 10.1016/j.immuni.2007.06.007. Publisher: Elsevier Ltd.

3. Patricia Barral, Paolo Polzella, Andreas Bruckbauer, Nico van Rooijen, Gurdyal S Besra, Vincenzo Cerundolo, and Facundo D Batista. >CD169+ macrophages present lipid antigens to mediate early activation of iNKT cells in lymph nodes. Nature Immunology, 11(4):303–312, April 2010. doi: 10.1038/ni.1853. Publisher: Nature Publishing Group.

4. Tri Giang Phan, Irina Grigorova, Takaharu Okada, and Jason G Cyster. Subcapsular encounter and complement-dependent transport of immune complexes by lymph node B cells. Nature Immunology, 8(9):992–1000, September 2007. doi: 10.1038/ni1494. Publisher: Nature Publishing Group.

5. Tobias Junt, E. Ashley Moseman, Matteo Iannacone, Steffen Massberg, Philipp A. Lang, Marianne Boes, Katja Fink, Sarah E. Henrickson, Dmitry M. Shayakhmetov, Nelson C. Di Paolo, Nico van Rooijen, Thorsten R. Mempel, Sean P. Whelan, and Ulrich H. von Andrian. Subcapsular sinus macrophages in lymph nodes clear lymph-borne viruses and present them to antiviral B cells. Nature, 450(7166):110–114, November 2007. ISSN 0028-0836, 1476-4687. doi: 10.1038/nature06287.

6. Tri Giang Phan, Jesse A Green, Elizabeth E Gray, Ying Xu, and Jason G Cyster. Immune complex relay by subcapsular sinus macrophages and noncognate B cells drives antibody affinity maturation. Nature Immunology, 10(7):786–793, June 2009. doi: 10.1038/ni.1745.

7. Federica Moalli, Steven T. Proulx, Reto Schwendener, Michael Detmar, Christoph Schlapbach, and Jens V. Stein. Intravital and Whole-Organ Imaging Reveals Capture of Melanoma-Derived Antigen by Lymph Node Subcapsular Macrophages Leading to Widespread Deposition on Follicular Dendritic Cells. Front. Immunol., 6, March 2015. ISSN 1664-3224. doi: 10.3389/fimmu.2015.00114.

8. Chung Park and John H Kehrl. An integrin/MFG-E8 shuttle loads HIV-1 viral-like particles onto follicular dendritic cells in mouse lymph node. eLife, 8:e47776, December 2019. ISSN 2050-084X. doi: 10.7554/eLife.47776.

9. Matteo Iannacone, E. Ashley Moseman, Elena Tonti, Lidia Bosurgi, Tobias Junt, Sarah E. Henrickson, Sean P. Whelan, Luca G. Guidotti, and Ulrich H. von Andrian. Subcapsular sinus macrophages prevent CNS invasion on peripheral infection with a neurotropic virus. Nature, 465(7301):1079–1083, June 2010. ISSN 0028-0836, 1476-4687. doi: 10.1038/nature09118.

10. Kenichi Asano, Ami Nabeyama, Yasunobu Miyake, Chun-Hong Qiu, Ai Kurita, Michio Tomura, Osami Kanagawa, Shin-ichiro Fujii, and Masato Tanaka. CD169-Positive Macrophages Dominate Antitumor Immunity by Crosspresenting Dead Cell-Associated Antigens. Immunity, 34(1):85–95, January 2011. ISSN 10747613. doi: 10.1016/j.immuni.2010.12.011.

11. Elodie Mohr, Karine Serre, Rudolf A. Manz, Adam F. Cunningham, Mahmood Khan, Deborah L. Hardie, Roger Bird, and Ian C. M. MacLennan. Dendritic Cells and Monocyte/Macrophages That Create the IL-6/APRIL-Rich Lymph Node Microenvironments Where Plasmablasts Mature. J Immunol, 182(4):2113–2123, February 2009. ISSN 0022-1767,1550-6606. doi: 10.4049/jimmunol.0802771.

12. Elizabeth E Gray and Jason G Cyster. Lymph node macrophages. J Innate Immun, 41(5-6):424–436, January 2012. doi: 10.1159/000337007. Publisher: Karger Publishers.

13. Katelyn M Spillane and Pavel Tolar. B cell antigen extraction is regulated by physical properties of antigen-presenting cells. Journal of Cell Biology, 216(1):217–230, January 2017. doi: 10.1083/jcb.201607064. Publisher: Rockefeller University Press.

14. Mohey Eldin El Shikh, Rania El Sayed, Andras K Szakal, and John G Tew. Follicular dendritic cell (FDC)-Fc*γ*RIIB engagement via immune complexes induces the activated FDC phenotype associated with secondary follicle development. European Journal of Immunology, 36(10):2715–2724, October 2006. doi: 10.1002/eji.200636122.

15. Balthasar A Heesters, Priyadarshini Chatterjee, Young-A Kim, Santiago F Gonzalez, Michael P Kuligowski, Tomas Kirchhausen, and Michael C Carroll. Endocytosis and Recycling of Immune Complexes by Follicular Dendritic Cells Enhances B Cell Antigen Binding and Activation. Immunity, 38(6):1164–1175, June 2013. doi: 10.1016/j.immuni.2013.02.023. Publisher: Elsevier Ltd.

16. Balthasar A. Heesters, Cees E. Van der Poel, and Michael C. Carroll. Follicular Dendritic Cell Isolation and Loading of Immune Complexes. In Dinis Pedro Calado, editor, Germinal Centers, volume 1623, pages 105–112. Springer New York, New York, NY, 2017. ISBN 978-1-4939-7094-0978-1-4939-7095-7. doi: 10.1007/978-1-4939-7095-7_9. Series Title: Methods in Molecular Biology.

17. Frederic Geissmann, Markus G. Manz, Steffen Jung, Michael H. Sieweke, Miriam Merad, and Klaus Ley. Development of Monocytes, Macrophages, and Dendritic Cells. Science, 327(5966):656–661, February 2010. ISSN 0036-8075, 1095-9203. doi: 10.1126/science.1178331.

18. Michael Y. Gerner, Parizad Torabi-Parizi, and Ronald N. Germain. Strategically Localized Dendritic Cells Promote Rapid T Cell Responses to Lymph-Borne Particulate Antigens. Immunity, 42(1):172–185, January 2015. ISSN 10747613. doi: 10.1016/j.immuni.2014.12.024.

19. Nagahito Saito, Karen A F Pulford, Jean-Marc Masse, and David Y Mason. Ultrastructural Localization of the CD68 Macrophage-associated Antigen in Human Blood Neutrophils and Monocytes. page 7.

20. Dante Alexander Patrick Louie and Shan Liao. Lymph Node Subcapsular Sinus Macrophages as the Frontline of Lymphatic Immune Defense. Front. Immunol., 10:347, February 2019. ISSN 1664-3224. doi: 10.3389/fimmu.2019.00347.

21. Abdouramane Camara, Alice C. Lavanant, Jun Abe, Henri Lee Desforges, Yannick O. Alexandre, Erika Girardi, Zinaida Igamberdieva, Kenichi Asano, Masato Tanaka, Thomas Hehlgans, Klaus Pfeffer, Sébastien Pfeffer, Scott N. Mueller, Jens V. Stein, and Christopher G. Mueller. CD169 ^+^macrophages in lymph node and spleen critically depend on dual RANK and LTbetaR signaling. Proc. Natl. Acad. Sci. U.S.A., 119(3):e2108540119, January 2022. ISSN 0027-8424, 1091-6490. doi: 10.1073/pnas.2108540119.

22. Danilo Pellin, Natalie Claudio, Zihan Guo, Tahereh Ziglari, and Ferdinando Pucci. Gene Expression Profiling of Lymph Node Sub-Capsular Sinus Macrophages in Cancer. Front. Immunol., 12:672123, June 2021. ISSN 1664-3224. doi: 10.3389/fimmu.2021.672123.

23. Valentin Jaumouillé, Yoav Farkash, Khuloud Jaqaman, Raibatak Das, Clifford A Lowell, and Sergio Grinstein. Actin cytoskeleton reorganization by Syk regulates Fc*γ*receptor responsiveness by increasing its lateral mobility and clustering. Developmental Cell, 29(5):534–546, June 2014. doi: 10.1016/j.devcel.2014.04.031. Publisher: Elsevier Inc.

24. Julia Riedl, Kevin C Flynn, Aurelia Raducanu, Florian Gärtner, Gisela Beck, Michael Bösl, Frank Bradke, Steffen Massberg, Attila Aszodi, Michael Sixt, and Roland Wedlich-Söldner. Lifeact mice for studying F-actin dynamics. Nature Methods, 7(3):168–169, March 2010. doi: 10.1038/nmeth0310-168. Publisher: Nature Publishing Group.

25. Joshua D Karslake, Eric D Donarski, Sarah A Shelby, Lucas M Demey, Victor J DiRita, Sarah L Veatch, and Julie S Biteen. SMAUG_ Analyzing single-molecule tracks with nonparametric Bayesian statistics. Methods, pages 1–11, April 2020. doi: 10.1016/j.ymeth.2020.03.008. Publisher: Academic Press.

26. Marina E Chicurel, Christopher S Chen, and Donald E Ingber. Cellular control lies in the balance of forces. Current Opinion in Cell Biology, 10(2):232–239, April 1998. ISSN 09550674. doi: 10.1016/S0955-0674(98)80145-2.

27. Tony Yeung, Penelope C. Georges, Lisa A. Flanagan, Beatrice Marg, Miguelina Ortiz, Makoto Funaki, Nastaran Zahir, Wenyu Ming, Valerie Weaver, and Paul A. Janmey. Effects of substrate stiffness on cell morphology, cytoskeletal structure, and adhesion. Cell Motil. Cytoskeleton, 60(1):24–34, January 2005. ISSN 0886-1544, 1097-0169. doi: 10.1002/cm.20041.

28. Bryant L. Doss, Meng Pan, Mukund Gupta, Gianluca Grenci, René-Marc Mège, Chwee Teck Lim, Michael P Sheetz, Raphael Voituriez, and Benoît Ladoux. Cell response to substrate rigidity is regulated by active and >passive cytoskeletal stress. Proc Natl Acad Sci USA, 117(23):12817–12825, June 2020. ISSN 0027-8424, 1091-6490. doi: 10.1073/pnas.1917555117.

29. Joan-Carles Escolano, Anna V. Taubenberger, Shada Abuhattum, Christine Schweitzer, Aleeza Farrukh, Aránzazu del Campo, Clare E. Bryant, and Jochen Guck. Compliant Substrates Enhance Macrophage Cytokine Release and NLRP3 Inflammasome Formation During Their Pro-Inflammatory Response. Front. Cell Dev. Biol., 9:639815, March 2021. ISSN 2296-634X. doi: 10.3389/fcell.2021.639815.

30. Hannah Schachtner, Simon D. J. Calaminus, Steven G. Thomas, and Laura M. Machesky. Podosomes in adhesion, migration, mechanosensing and matrix remodeling: Podosomes. Cytoskeleton, 70(10):572–589, October 2013. ISSN 19493584. doi: 10.1002/cm.21119.

31. Madison Bolger-Munro, K Choi, J M Scurll, A Libin, R S Chappell, Duke Sheen, May Dang-Lawson, Xufeng Wu, John J Priatel, Daniel Coombs, John A Hammer III, and Michael R Gold. Arp2/3 complex-driven spatial patterning of the BCR enhances immune synapse formation, BCR signaling and B cell activation. Elife, 8:e44574, 2019. doi: 10.7554/eLife.44574.001.

32. Sophie I Roper, Laabiah Wasim, Dessislava Malinova, Michael Way, Susan Cox, and Pavel Tolar. B cells extract antigens at Arp2/3-generated actin foci interspersed with linear filaments. eLife, 8:e48093, December 2019. ISSN 2050-084X. doi: 10.7554/eLife.48093.

33. Deniz Saltukoglu, Bugra Özdemir, Michael Holtmannspötter, Ralf Reski, Jacob Piehler, Rainer Kurre, and Michael Reth. Plasma membrane topography governs the threedimensional dynamic localization of IgM B cell receptor clusters. bioRxiv, April 2022. doi: 10.1101/2022.04.29.489661.

34. En Cai, Kyle Marchuk, Peter Beemiller, Casey Beppler, Matthew G Rubashkin, Valerie M Weaver, Audrey Gérard, Tsung-Li Liu, Bi-Chang Chen, Eric Betzig, Frederic Bartumeus, and Matthew F Krummel. Visualizing dynamic microvillar search and stabilization during ligand detection by T cells. Science, 356(6338):eaal3118, May 2017. doi: 10.1126/science.aal3118. Publisher: American Association for the Advancement of Science.

35. Jia C Wang, Yang-In Yim, Xufeng Wu, Valentin Jaumouille, Andrew Cameron, Clare M Waterman, John H Kehrl, and John A Hammer. A B-cell actomyosin arc network couples integrin co-stimulation to mechanical force-dependent immune synapse formation. eLife, 11:e72805, April 2022. ISSN 2050-084X. doi: 10.7554/eLife.72805.

36. Elizabeth Natkanski, Wing-Yiu Lee, Bhakti Mistry, Antonio Casal, Justin E Molloy, and Pavel Tolar. B Cells Use Mechanical Energy to Discriminate Antigen Affinities. Science, 340 (6140):1587–1590, June 2013. doi: 10.1126/science.1237572. Publisher: American Association for the Advancement of Science.

37. Katelyn M Spillane and Pavel Tolar. Mechanics of antigen extraction in the B cell synapse. Molecular Immunology, 101:319–328, July 2018. doi: 10.1016/j.molimm.2018.07.018. Publisher: Elsevier.

38. Mark Bennett, Marco Cantini, Julien Reboud, Jonathan M Cooper, Pere Roca-Cusachs, and Manuel Salmeron-Sanchez. Molecular clutch drives cell response to surface viscosity. Proc. Natl.Acad. Sci. U.S.A., 115(6):1192–1197, February 2018. doi: 10.1073/pnas.1710653115.

39. Christina Ketchum, Heather Miller, Wenxia Song, and Arpita Upadhyaya. Ligand Mobility Regulates B Cell Receptor Clustering and Signaling Activation. Biophysical Journal, 106 (1):26–36, January 2014. doi: 10.1016/j.bpj.2013.10.043. Publisher: Cell Press.

40. Bebhinn Treanor, David Depoil, Aitor Gonzalez-Granja, Patricia Barral, Michele Weber, Omer Dushek, Andreas Bruckbauer, and Facundo D Batista. The Membrane Skeleton Controls Diffusion Dynamics and Signaling through the B Cell Receptor. Immunity, 32(2): 187-199, February 2010. doi: 10.1016/j.immuni.2009.12.005. Publisher: Elsevier Ltd.

41. Pavel Tolar, Joseph Hanna, Peter D Krueger, and Susan K Pierce. The Constant Region of the Membrane Immunoglobulin Mediates B Cell-Receptor Clustering and Signaling in Response to Membrane Antigens. Immunity, 30(1):44–55, January 2009. doi: 10.1016/j.immuni.2008.11.007. Publisher: Elsevier Ltd.

42. Carla R Nowosad, Katelyn M Spillane, and Pavel Tolar. Germinal center B cells recognize antigen through a specialized immune synapse architecture. Nature Immunology, 17(7):870–877, January 2016. doi: 10.1038/ni.3458. Publisher: Nature Publishing Group.

43. Charlotte M. Fonta, Simon Arnoldini, Daniela Jaramillo, Alessandra Moscaroli, Annette Oxenius, Martin Behe, and Viola Vogel. Fibronectin fibers are highly tensed in healthy organs in contrast to tumors and virus-infected lymph nodes. Matrix Biology Plus, 8:100046, November 2020. ISSN 25900285. doi: 10.1016/j.mbplus.2020.100046.

44. Harry L. Horsnell, Robert J. Tetley, Henry De Belly, Spyridon Makris, Lindsey J. Millward, Agnesska C. Benjamin, Lucas A. Heeringa, Charlotte M. de Winde, Ewa K. Paluch, Yanlan Mao, and Sophie E. Acton. Lymph node homeostasis and adaptation to immune challenge resolved by fibroblast network mechanics. Nat Immunol, 23(8):1169–1182, August 2022. ISSN 1529-2908, 1529-2916. doi: 10.1038/s41590-022-01272-5.

45. Frank P. Assen, Jun Abe, Miroslav Hons, Robert Hauschild, Shayan Shamipour, Walter A. Kaufmann, Tommaso Costanzo, Gabriel Krens, Markus Brown, Burkhard Ludewig, Simon Hippenmeyer, Carl-Philipp Heisenberg, Wolfgang Weninger, Edouard Hannezo, Sanjiv A. Luther, Jens V. Stein, and Michael Sixt. Multitier mechanics control stromal adaptations in the swelling lymph node. Nat Immunol, 23(8):1246–1255, August 2022. ISSN 1529-2908, 1529-2916. doi: 10.1038/s41590-022-01257-4.

46. W. E. Allen, G. E. Jones, J. W. Pollard, and A. J. Ridley. Rho, Rac and Cdc42 regulate actin organization and cell adhesion in macrophages. J Cell Sci, 110 (Pt 6):707–720, March 1997. ISSN 0021-9533. doi: 10.1242/jcs.110.6.707.

47. A. J. Ridley, H. F. Paterson, C. L. Johnston, D. Diekmann, and A. Hall. The small GTP-binding protein rac regulates growth factor-induced membrane ruffling. Cell, 70(3):401–410, August 1992. ISSN 0092-8674. doi: 10.1016/0092-8674(92)90164-8.

48. R Kozma, S Ahmed, A Best, and L Lim. The Ras-related protein Cdc42Hs and bradykinin promote formation of peripheral actin microspikes and filopodia in Swiss 3T3 fibroblasts. Mol Cell Biol, 15(4):1942–1952, April 1995. ISSN 0270-7306, 1098-5549. doi: 10.1128/MCB.15.4.1942.

49. Alberto Elosegui-Artola, Elsa Bazellières, Michael D Allen, Ion Andreu, Roger Oria, Raimon Sunyer, Jennifer J Gomm, John F Marshall, J Louise Jones, Xavier Trepat, and Pere Roca-Cusachs. Rigidity sensing and adaptation through regulation of integrin types. Nature Materials, 13(6):631–637, June 2014. doi: 10.1038/nmat3960. Publisher: Nature Publishing Group.

50. Leontios Pappas, Mathilde Foglierini, Luca Piccoli, Nicole L. Kallewaard, Filippo Turrini, Chiara Silacci, Blanca Fernandez-Rodriguez, Gloria Agatic, Isabella Giacchetto-Sasselli, Gabriele Pellicciotta, Federica Sallusto, Qing Zhu, Elisa Vicenzi, Davide Corti, and Antonio Lanzavecchia. Rapid development of broadly influenza neutralizing antibodies through re-dundant mutations. Nature, 516(7531):418–422, December 2014. ISSN 0028-0836,14764687. doi: 10.1038/nature13764.

51. Florian Klein, Ron Diskin, Johannes F. Scheid, Christian Gaebler, Hugo Mouquet, Ivelin S. Georgiev, Marie Pancera, Tongqing Zhou, Reha-Baris Incesu, Brooks Zhongzheng Fu, Priyanthi N.P. Gnanapragasam, Thiago Y. Oliveira, Michael S. Seaman, Peter D. Kwong, Pamela J. Bjorkman, and Michel C. Nussenzweig. Somatic Mutations of the Immunoglobulin Framework Are Generally Required for Broad and Potent HIV-1 Neutralization. Cell, 153 (1):126–138, March 2013. ISSN 00928674. doi: 10.1016/j.cell.2013.03.018.

52. Sophie E. Acton, Aaron J. Farrugia, Jillian L. Astarita, Diego Mourão-Sá, Robert P. Jenkins, Emma Nye, Steven Hooper, Janneke van Blijswijk, Neil C. Rogers, Kathryn J. Snelgrove, Ian Rosewell, Luis F. Moita, Gordon Stamp, Shannon J. Turley, Erik Sahai, and Caetano Reis e Sousa. Dendritic cells control fibroblastic reticular network tension and lymph node expansion. Nature, 514(7523):498–502, October 2014. ISSN 0028-0836, 1476-4687. doi: 10.1038/nature13814.

53. Johannes Schindelin, Ignacio Arganda-Carreras, Erwin Frise, Verena Kaynig, Mark Longair, Tobias Pietzsch, Stephan Preibisch, Curtis Rueden, Stephan Saalfeld, Benjamin Schmid, Jean-Yves Tinevez, Daniel James White, Volker Hartenstein, Kevin Eliceiri, Pavel Tomancak, and Albert Cardona. Fiji: an open-source platform for biological-image analysis. Nat Methods, 9(7):676–682, July 2012. ISSN 1548-7091,1548-7105. doi: 10.1038/nmeth.2019.

54. Fabrice de Chaumont, Stéphane Dallongeville, Nicolas Chenouard, Nicolas Hervé, Sorin Pop, Thomas Provoost, Vannary Meas-Yedid, Praveen Pankajakshan, Timothée Lecomte, Yoann Le Montagner, Thibault Lagache, Alexandre Dufour, and Jean-Christophe Olivo-Marin. Icy: an open bioimage informatics platform for extended reproducible research. Nat Methods, 9(7):690–696, July 2012. ISSN 1548-7091,1548-7105. doi: 10.1038/nmeth.2075.

55. Samuel J. Lord, Katrina B. Velle, R. Dyche Mullins, and Lillian K. Fritz-Laylin. SuperPlots: Communicating reproducibility and variability in cell biology. Journal of Cell Biology, 219(6):e202001064, June 2020. ISSN 0021-9525, 1540-8140. doi: 10.1083/jcb.202001064.

56. Jeffrey L. Hutter and John Bechhoefer. Calibration of atomic-force microscope tips. Review of Scientific Instruments, 64(7):1868–1873, July 1993. ISSN 0034-6748, 1089-7623. doi: 10.1063/1.1143970.

57. Félix Rico, Pere Roca-Cusachs, Núria Gavara, Ramon Farré, Mar Rotger, and Daniel Navajas. Probing mechanical properties of living cells by atomic force microscopy with blunted pyramidal cantilever tips. Phys. Rev. E, 72(2):021914, August 2005. ISSN 1539-3755, 1550-2376. doi: 10.1103/PhysRevE.72.021914.

58. M.R. Bubb, A.M. Senderowicz, E.A. Sausville, K.L. Duncan, and E.D. Korn. Jasplakinolide, a cytotoxic natural product, induces actin polymerization and competitively inhibits the binding of phalloidin to F-actin. Journal of Biological Chemistry, 269(21):14869–14871, May 1994. ISSN 00219258. doi: 10.1016/S0021-9258(17)36545-6.

59. Xiaoyu Sun, Donovan Y.Z. Phua, Lucas Axiotakis, Mark A. Smith, Elizabeth Blankman, Rui Gong, Robert C. Cail, Santiago Espinosa de los Reyes, Mary C. Beckerle, Clare M. Waterman, and Gregory M. Alushin. Mechanosensing through Direct Binding of Tensed F-Actin by LIM Domains. Developmental Cell, 55(4):468–482.e7, November 2020. ISSN 15345807. doi: 10.1016/j.devcel.2020.09.022.

60. Kota Miura. Bleach correction ImageJ plugin for compensating the photobleaching of timelapse sequences. F1000Res, 9:1494, December 2020. ISSN 2046-1402. doi: 10.12688/f1000research.27171.1.

61. Jean-Yves Tinevez, Nick Perry, Johannes Schindelin, Genevieve M. Hoopes, Gregory D. Reynolds, Emmanuel Laplantine, Sebastian Y. Bednarek, Spencer L. Shorte, and Kevin W. Eliceiri. TrackMate: An open and extensible platform for single-particle tracking. Methods, 115:80–90, February 2017. ISSN 10462023. doi: 10.1016/j.ymeth.2016.09.016.

62. Katelyn M Spillane, Jaime Ortega-Arroyo, Gabrielle de Wit, Christian Eggeling, Helge Ewers, Mark I Wallace, and Philipp Kukura. High-Speed Single-Particle Tracking of GM1 in Model Membranes Reveals Anomalous Diffusion due to Interleaflet Coupling and Molecular Pinning. Nano Letters, 14(9):5390–5397, September 2014. doi: 10.1021/nl502536u. Publisher: American Chemical Society.

